# Natural variation of warm temperature-induced raffinose accumulation identifies *TREHALOSE-6-PHOSPHATE SYNTHASE 1* as a modulator of thermotolerance

**DOI:** 10.1101/2023.05.15.540763

**Authors:** Niklas Reichelt, Arthur Korte, Markus Krischke, Martin J. Mueller, Daniel Maag

## Abstract

High temperature stress limits plant growth and reproduction. Exposure to high temperature, however, also elicits a conserved physiological response, which protects plants from the damage evoked by heat. This response involves a partial reconfiguration of the plant metabolome including the accumulation of the trisaccharide raffinose. In this study, we explored the intra-specific variation of warm temperature-induced raffinose accumulation as a metabolic marker for temperature responsiveness with the aim to identify genes that contribute to plant thermotolerance. By combining raffinose measurements in 250 *Arabidopsis thaliana* accessions following a mild heat treatment with genome-wide association studies we identified five genomic regions that were associated with the observed trait variation. Subsequent functional analyses confirmed a causal relationship between *TREHALOSE-6-PHOSPHATE SYNTHASE 1* (*TPS1*) and warm temperature-dependent raffinose synthesis. Moreover, complementation of the *tps1-1* null mutant with functionally distinct TPS1 isoforms differentially affected carbohydrate metabolism under more severe heat stress. While higher TPS1 activity was associated with reduced endogenous sucrose levels and thermotolerance, disruption of trehalose 6-phosphate signalling resulted in higher accumulation of transitory starch and sucrose and was associated with enhanced heat resistance. Taken together, our findings suggest a role of trehalose 6-phosphate in thermotolerance most likely through its regulatory function in carbon partitioning and sucrose homeostasis.

## Introduction

Temperature is a key determinant of plant growth and performance. Accordingly, extreme temperatures can exert detrimental effects on a plant’s reproductive success, or in the case of crop plants, yield (Battisti and Naylor, 2009). At the cellular level, severe heat stress leads to the over-accumulation of reactive oxygen species, membrane destabilisation and protein unfolding (Mittler *et al*., 2012). At the same time exposure to high temperatures causes a decline in photosynthetic rates that is accompanied by enhanced photorespiration (Moore *et al*., 2021). As a consequence of the overall reduction in carbon assimilation, the formation of transitory starch is largely compromised by heat stress (Thalmann and Santelia, 2017).

However, plants are able to adapt to adverse temperature conditions. Likewise, increasing temperatures elicit a conserved physiological response, which mitigates the damage evoked by heat and which is indispensable for the survival of heat stress (Schöffl *et al*., 1998, Mittler *et al*., 2012). This heat shock response (HSR) includes the rapid accumulation of HEAT SHOCK PROTEINS (HSPs), many of which function as molecular chaperones that are involved in protein homeostasis. During heat stress, these HSPs are involved in preventing aggregation of denatured proteins by keeping them in a folding-competent state or in facilitating their degradation (Waters and Vierling, 2020, Guihur *et al*., 2022). The majority of the heat-responsive *HSP* genes is under positive transcriptional regulation by specific HEAT SHOCK TRANSCRIPTION FACTORS (HSFs). The genome of Arabidopsis (*Arabidopsis thaliana* (L.) Heynh.) encodes 21 HSFs divided into three distinct classes (A, B and C). Out of this regulatory network three functionally redundant homologues (HSFA1a, b and d) have been identified as the master regulators of the HSR (Liu *et al*., 2011). However, additional signalling pathways were found to contribute to the acquisition of thermotolerance and this contribution is partly independent of the accumulation of HSPs (Larkindale *et al*., 2005, Li *et al*., 2019). In fact, *HSPs* only constitute a small portion of the heat-responsive transcriptome in Arabidopsis (Larkindale and Vierling, 2008). Correspondingly, exposure to high temperature also induces the expression of genes coding for proteins involved in oxidative stress regulation and other metabolic processes, and leads to a partial reconfiguration of the plant metabolome including the accumulation of raffinose-family oligosaccharides (RFOs) (Panikulangara *et al*., 2004, Larkindale and Vierling, 2008, Mueller *et al*., 2015, Mueller *et al*., 2017).

The first committed step within the RFO biosynthetic pathway, i.e., the synthesis of galactinol from UDP-galactose and *myo*-inositol, is transcriptionally regulated in a temperature-dependent manner *via* binding of HSFA1 to the promotor region of *GALACTINOL SYNTHASE 1* (*GOLS1*) (Panikulangara *et al*., 2004, Mueller *et al*., 2015, Cortijo *et al*., 2017). The predominant RFO in Arabidopsis is raffinose. Under high temperature conditions, raffinose is synthesised by RAFFINOSE SYNTHASE 5 (RS5) *via* transfer of the galactose moiety from galactinol onto a molecule of sucrose (Egert *et al*., 2013). Curiously, *RS5* expression is downregulated at elevated temperatures (*GENEVESTIGATOR*, Version 9.6.1; Hruz *et al*., 2008). Although RFO-deficient *gols1* mutant plants are not impaired in thermotolerance (Panikulangara *et al.,* 2004) recent publications suggest a beneficial role for RFOs in heat stress resistance (Yan *et al*., 2022). Accordingly, overexpression of *GOLS1* homologs from other plant species in Arabidopsis not only results in an increased accumulation of RFOs but also in enhanced thermotolerance, most likely due to increased scavenging of reactive oxygen species and a concomitant reduction in lipid peroxidation (Gu *et al*., 2016, Salvi *et al*., 2018, Gu *et al*., 2019). Taken together, these observations suggest that RFOs, including raffinose itself, can be considered as metabolic markers indicative of both HSFA1 activity at the molecular level and heat tolerance at the level of the whole plant.

Arabidopsis is widely distributed across the northern hemisphere with dispersed populations showing signatures of local adaptation to their natural habitats (Fournier-Level *et al*., 2011, Clauw *et al*., 2022). As a consequence, Arabidopsis displays a high degree of intraspecific variation including many stress-related traits. Recently, the genomes of 1135 Arabidopsis accessions have been sequenced and made publicly available by the 1001 Genomes Project (Alonso-Blanco *et al*., 2016) thereby providing a powerful resource for genome-wide association studies (GWAS). This resource has been used to identify genes that are involved in various abiotic stress responses such as salt (Awlia *et al*., 2021), submergence (Meng *et al*., 2022) and nutrient stress (Bouain *et al*., 2019). Although recent studies suggest that Arabidopsis also shows a high degree of intraspecific variation concerning a variety of responses towards high temperatures (Tonsor *et al*., 2008, Barah *et al*., 2013, Wolfe and Tonsor, 2014, Scheepens *et al*., 2018), this variation has hardly been used to identify the underlying regulatory genes.

In this study, we identify *TREHALOSE-6-PHOSPHATE SYNTHASE 1* (*TPS1*; AT1G78580) as a modulator of thermotolerance in Arabidopsis by exploiting the natural variation of warm temperature-induced raffinose accumulation. Following the phenotyping of 250 Arabidopsis accessions and subsequent genome-wide association analyses, we first confirm the causal relationship between TPS1 activity and the accumulation of RFOs at high temperatures using different *tps1* TILLING lines. We then demonstrate that complementation of the *tps1-1* null mutant with functionally distinct TPS1 isoforms leads to extensive changes in carbohydrate metabolism under heat stress and differentially affects the accumulation of soluble sugars and transitory starch under these conditions. Finally, we show that the increased accumulation of transitory starch in *TPS1* complementation lines with impaired T6P signalling during heat stress is accompanied by enhanced long-term thermotolerance while complementation with a more active TPS1 isoform is associated with reduced heat resistance.

## Materials & Methods

### Plant material

The GWA mapping panel used in this study consisted of 250 natural accessions of *A. thaliana* (Extended Data 1) from the 1001 Genomes Project (Alonso-Blanco *et al*., 2016). Seeds were kindly provided by Magnus Nordborg (Gregor Mendel Institute, Vienna, Austria) and propagated for one generation in our greenhouse prior to the large-scale phenotyping. Seeds of the *tps1-11* and *tps1-12* TILLING lines (Gómez *et al*., 2010) as well as *rhm1-2* (also known as *rol1-2*) (Diet *et al*., 2006) were obtained from the Nottingham Arabidopsis Stock Centre (NASC), tested for the presence of the described mutations *via* sequencing, and propagated for at least one generation together with the corresponding wild-type plants in temperature-controlled climate chambers. The *TPS1* complementation lines *TPS1[ΔN]* and *TPS1[ΔNΔC]* (Fichtner *et al*., 2020) were kindly provided by John Lunn (Max Planck Institute of Molecular Plant Physiology, Potsdam-Golm, Germany).

### Plant growth and phenotyping conditions

For the large-scale phenotyping and the experiments with *tps1-11*, *tps1-12* and *rhm1-2* under the phenotyping conditions, seeds were sown on agar plates with ½-strength Murashige-Skoog (MS) medium (Duchefa Biochemie B.V., Haarlem, Netherlands), pH 5.7, containing 3% (w/v) sucrose and 1.2% (w/v) phyto agar (Duchefa Biochemie B.V.), and stratified for 7 days at 4 °C in the dark. After stratification plates were transferred to a growth chamber with 9 h/15 h (day/night) photoperiod set to 22 °C and 100-120 µmol m^-2^ s^-1^ (white fluorescent light) in a randomised manner and regularly shifted within the growth chamber to avoid any positional effects on seedling establishment and growth. On day 15, seedlings were transferred to a pre-heated growth chamber at 32 °C at Zeitgeber Time (ZT) 2, i.e., two hours after onset of the light period. Control seedlings remained in the 22 °C growth chamber. After four hours seedlings were harvested, flash-frozen in liquid nitrogen and stored at –80 °C until extraction. Due to spatial constraints, 17 accessions were treated on a given day and the phenotyping was thus distributed across 18 experimental days. To control for potential day-to-day variation concerning the 32 °C treatment each batch of 17 accessions contained Col-0 and Furni-1, to serve as internal controls for the heat treatment.

For experiments with the different *TPS1* complementation lines, seeds were sown on ½-strength MS plates, pH 5.7, containing 1% (w/v) sucrose and 1.2% (w/v) phyto agar, and stratified for 48 hours at 4 °C in the dark. After ten days of growth under the conditions described above, seedlings were transferred to new plates without sugar supplementation. This growing scheme was selected since under our experimental conditions *TPS1[ΔN]* showed a severe growth retardation phenotype that could be rescued by supplementing the growth medium with 1% sucrose (Figure S1). However, since sugar supplementation might also mask other *TPS1*-related phenotypes we decided to grow all seedlings on 1% sucrose-supplemented medium for ten days before they were transferred to sucrose-free medium and then grown for another four days. This resulted in seedlings that were similar in size at the beginning of the heat treatments but were not supplemented with an external carbon source (Figure S1).

### Extraction and quantification of soluble carbohydrates and starch

Soluble carbohydrates were extracted as described by Mueller *et al*. (2015) with modifications. In brief, approximately 25 mg of frozen plant material were extracted with 1 mL of chloroform/methanol (3:2, v/v; 0.06% (w/v) butylated hydroxytoluene, w/v) using a bead mill at 21 Hz for 10 minutes (Retsch, Haan, Germany). Subsequently, 200 µL of water were added to the samples followed by agitation at 21 Hz for another 10 minutes. Following centrifugation at 20 800 x *g* and 4 °C for 10 minutes, 450 µL of the upper, aqueous phase were transferred to a fresh reaction tube and evaporated to dryness using a rotational evaporator. The residual pellet containing the soluble carbohydrates was resuspended in 30 µL of 50% (v/v) methanol and stored at –20 °C until analysis by UPLC-MS/MS.

For the simultaneous extraction and quantification of soluble carbohydrates and starch from a single sample, approximately 20 mg of frozen plant material were ground to a fine powder and extracted three times by adding 1 mL of 90% (v/v) ethanol followed by an incubation at 60 °C for 5 minutes and centrifugation at 15,000 x *g* and 4 °C for 3 minutes. The first two fractions were recovered and combined, evaporated to dryness using a rotational evaporator and resuspended in 30 µl of 50% methanol (v/v) for quantification of soluble carbohydrates by UPLC-MS/MS. The third fraction was discarded. The residual pellet containing the insoluble starch was resuspended in 500 µL of 0.5 M KOH and incubated at 100 °C for 5 minutes to gelatinise the starch. Subsequently, the pH was adjusted to 4.8 by adding 350 µL of 0.4 M H_3_PO_4_. Following the addition of 100 µL of an enzyme mix containing α-amylase (0.5 U, Sigma-Aldrich, St. Louis, MO USA; A3176) and amyloglucosidase (7 U, Sigma-Aldrich; 10115) in 50 mM phosphate buffer (pH 7.4), samples were incubated for one hour at room temperature to allow for enzymatic hydrolysis of the starch. Following centrifugation, the supernatants containing the resultant glucose moieties were used for the fluorometric quantification assay. The reaction mixture consisted of 0.5 µL 10 mM Amplex Red (Invitrogen, Waltham, MA, USA; A22188), 1 µL horseradish peroxidase (0.01 U, Sigma-Aldrich; 77332), 1 µL glucose oxidase (0.1 U, Sigma-Aldrich; G7141), 47.5 µL 100 mM phosphate buffer (pH 7.4) and 50 µL of a given sample diluted 1:10 or 1:20 in 100 mM phosphate buffer (pH 7.4). Fluorescence was detected after a 30-minute incubation at room temperature in the dark using a Fluoroskan Ascent microplate reader (Thermo Fisher Scientific, Waltham, USA) at an excitation wavelength of 530 nm and an emission wavelength of 590 nm. Quantification was based on a calibration curve obtained from freshly hydrolysed pure starch (Sigma-Aldrich; 85642) at four different concentrations.

Soluble carbohydrates were detected using a Waters Acquity UPLC system coupled to a Waters Micromass Quattro Premier triple quadrupole mass spectrometer (Waters, Milford, CT, USA) operated in multiple reaction monitoring (MRM) mode. Chromatographic separation was achieved using an Acquity UPLC BEH Amide column (2.1 x 100 mm, 1.7 µm particle size) maintained at 40 °C with a linear binary gradient of 10-55% eluent B over 7.5 minutes at a flow rate of 0.2 mL/min. Eluent A consisted of 80% (v/v) acetonitrile in water (containing 0.1% (v/v) ammonium hydroxide) and eluent B consisted of 30% (v/v) acetonitrile in water (containing 0.1% (v/v) ammonium hydroxide). After chromatographic separation soluble carbohydrates were detected by MS coupled with an electrospray ionisation (ESI) source operated in negative mode as described by Mueller at al. (2015). The MRM transitions that were monitored are given in Table S1. For quantification 2 µg of α,α-trehalose-1,1‘-d_2_ (Sigma-Aldrich; 700452) and 8 µg of D-glucose-6,6-d_2_ (Sigma-Aldrich; 282650) were added to each sample as internal standards at the beginning of the extraction procedure and experimentally determined response factors were used for each analyte/standard pair (Table S1).

### Genome-wide association mapping

Genome-wide association mapping was performed separately on raffinose concentration data from plants grown at 22 °C and from plants exposed to 32 °C for 4 hours. The phenotypic data will be made available in the AraPheno database (Togninalli *et al*., 2020). An imputed version of the genotypic data for the 250 accessions was used (Arouisse *et al*., 2020). The association analysis was performed with permGWAS (John *et al*., 2022) and was based on a linear mixed model that corrected for population structure. Only SNPs with a minor allele frequency above 0.05 were considered, resulting in 1,420,453 SNPs for the final model. Based on these SNPs, the Bonferroni-corrected significance threshold was calculated to be 3.52 x 10^-8^. Manhattan and local plots were generated in *R* and the respective scripts are available at https://github.com/arthurkorte/GWAS.

### Survival assays

Tolerance towards long-term heat stress was assessed by exposing 14-day-old seedlings to a heat treatment at 37 °C for 48 hours. Following a recovery period of 14 days under control growth conditions, seedling survival was determined as the fraction of seedlings within a given agar plate that had grown new leaves. To account for any positional effects during the heat treatment, mutant lines were grown together with their corresponding wild type on replicate plates.

### Statistical analysis

Metabolite and survival data were analysed by one-or multi-factorial analysis of variance (ANOVA) followed by a Tukey HSD test. Normality of the data was assessed based on the residuals of the ANOVA model and using the *plotresid* function of the *R* package *RVAideMemoire* (Version 0.9-80) (Hervé, 2015). Variance homogeneity was tested using Levene’s test within the R package *car* (Version 3.1-0) (Fox and Weisberg, 2019). If necessary, data were log10-transformed to meet the requirements for ANOVA. All analyses were conducted in *R* (Version 4.1.1).

## Results

### *Arabidopsis* shows extensive natural variation for warm temperature-induced raffinose accumulation

In order to identify genes that contribute to warm temperature signalling in *Arabidopsis* we measured the accumulation of the trisaccharide raffinose in 250 accessions from the 1001 Genomes Project (Alonso-Blanco *et al*., 2016) after a mild heat treatment, i.e. a four-hour exposure to 32 °C, as a marker for temperature responsiveness. Under our experimental conditions, this was the minimum temperature that was required to induce a significant increase in raffinose levels in Columbia-0 (Col-0) and was thus considered a well-suited treatment for the detection of intra-specific trait variation (Figure S2). Moreover, 32 °C also defined the minimum acclimation temperature that was needed for the acquisition of short-term thermotolerance to a more severe heat stress in Col-0, and coincided with the temperature range at which substantial genotypic variation of short-term acquired thermotolerance was observed across 25 natural accessions (Figure S2).

Quantification of raffinose levels within the mapping population revealed a high degree of intra-specific variation with concentrations ranging from 2.5 nmol/g FW to 231 nmol/g FW and 24.9 nmol/g FW to 682 nmol/g FW under control and warm temperature conditions, respectively (Figure 1a and Extended Data 1). While we observed a strong positive correlation between raffinose levels at 22 °C and 32 °C (*r* = 0.693, *p* < 0.001, Spearman’s rank correlation test, Figure S3), only a weak, yet statistically significant, positive correlation was observed between raffinose levels at either temperature and latitude of origin of the 250 accessions (*r* = 0.222, *p* < 0.001 for 22 °C and *r* = 0.285, *p* < 0.001 for 32 °C, Spearman’s rank correlation test) indicating that accessions from colder climates contained more raffinose irrespectively of the temperature treatment (Figure S3). Therefore, we assessed next whether raffinose levels were correlated with any of the temperature-related bioclimatic variables from the WorldClim database (Fick and Hijmans, 2017). Indeed, raffinose concentrations at both experimental temperatures were negatively correlated with annual mean temperature, maximum temperature of the warmest month and the mean temperature of the warmest quarter (Figure S4).

**Figure 1.**
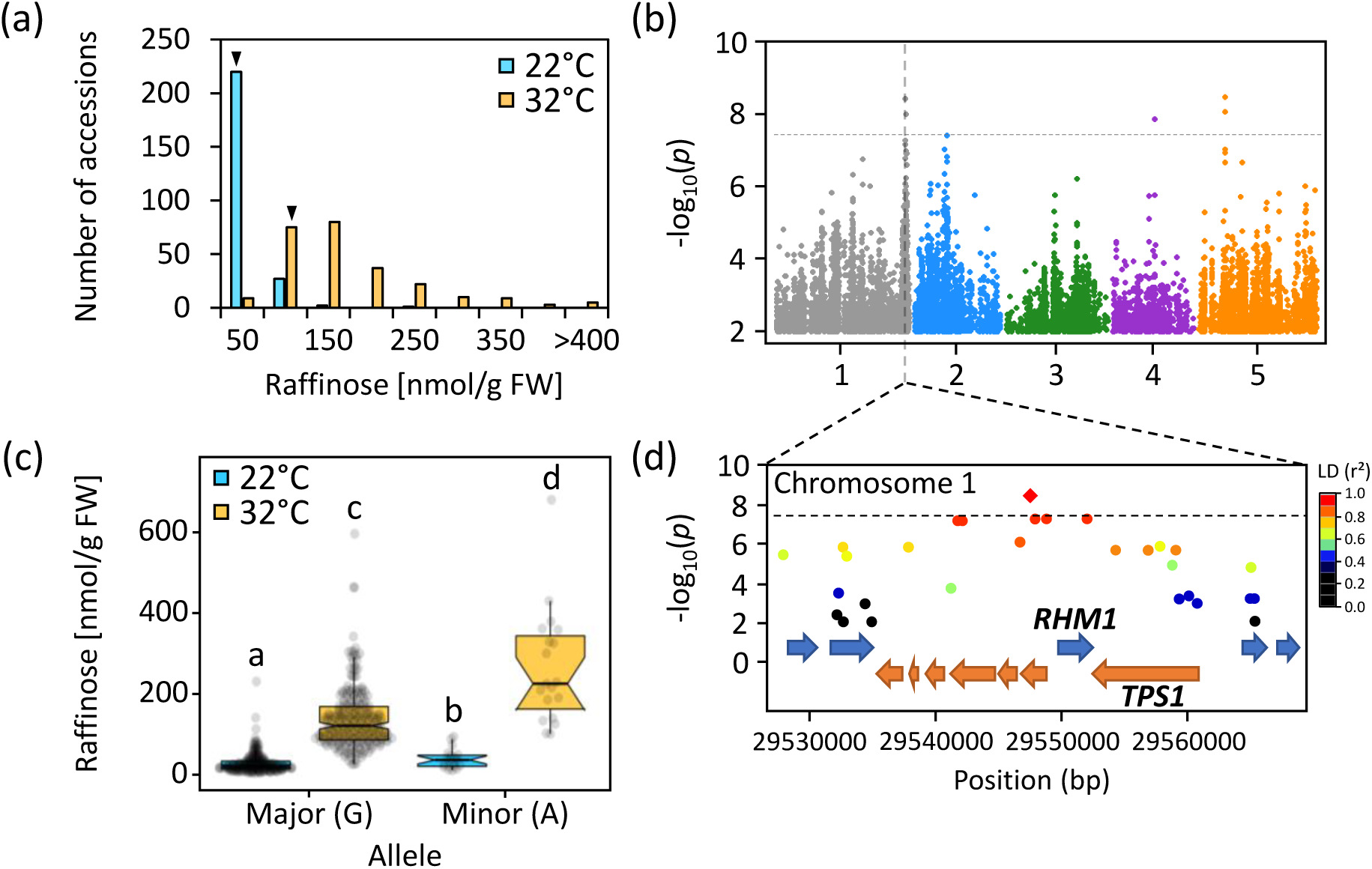
Natural variation of warm temperature-induced raffinose accumulation is associated with a genetic locus at the end of chromosome 1. (a) Distribution of raffinose concentrations in 14-day-old seedlings of 250 *Arabidopsis thaliana* accessions that were exposed to 32 °C for 4 hours (orange) or maintained at 22 °C as control (blue). Arrows indicate the relative position of Columbia-0 at the different temperatures. *N* = 3. (b) Manhattan plot depicting the statistical association between individual single-nucleotide polymorphisms (SNPs) and the variation in raffinose levels at 32 °C on a –log_10_(*p*) scale. The dashed line indicates the Bonferroni-corrected significance threshold. (c) Raffinose concentrations for accessions grouped according to their allelic state in position 29,547,498 on chromosome 1. At this position 231 accessions carried the major allele (guanine) while 19 accessions carried the minor allele (adenine). Box plots display the first and third quartiles with the median as centre. Notches indicate 95% confidence intervals. Different letters above the boxes indicate statistically significant differences (two-way ANOVA followed by Tukey’s HSD test, *p* < 0.05). (d) Local plot of the genetic locus around the most significant SNP in position 29,547,498 on chromosome 1 (diamond symbol). The linkage disequilibrium (LD) relative to this SNP is depicted on a colour scale (r²). Open reading frames are depicted in their respective orientation along the chromosome. The dashed line indicates the Bonferroni-corrected significance threshold of the genome-wide association study.

### Genome-wide association mapping identifies candidate genes that are associated with variation in temperature-dependent raffinose accumulation

To identify genomic regions that were associated with the observed variation in raffinose levels at 22 °C and 32 °C, two separate GWAS analyses were performed using a linear mixed model approach (see Figures S5 and S6 for all Manhattan and QQ-Plots). Concerning the warm temperature treatment five statistically significant SNPs were detected applying a Bonferroni-corrected significance threshold (Figure 1b and Table 1). These SNPs were located on chromosomes 1 (2 SNPs), 4, and 5 (2 SNPs). Genes within a 5-kb window to either side of the significant SNPs were considered candidate genes and are listed in Table S2. This list was narrowed down by removing genes that were barely expressed in Col-0 seedlings that were grown under similar conditions, based on an in-house transcriptomic data set (Table S1), resulting in the identification of 13 candidate genes (Table 1).

**Table 1.** Single-nucleotide polymorphisms that were statistically significantly associated with the variation of warm temperature-induced raffinose accumulation at 32 °C and the underlying candidate genes.

| *SNP<sup>a</sup>* | *p-value* | *Associated gene<sup>b</sup>* | *Name* | *Position on chromosome* | *Regulation<sup>c</sup>* |
| --- | --- | --- | --- | --- | --- |
| Chr 1, Pos 29,547,498 | 3.51 x 10<sup>-9</sup> | AT1G78540 | *SHB* | 29,540,983 – 29,544,875 | -- |
|  |  | AT1G78550 | na | 29,544,901 – 29,546,511 | -- |
|  |  | AT1G78560 | *BASS1* | 29,546,558 – 29,548,866 | -- |
|  |  | AT1G78570 | *RHM1* | 29,549,539 – 29,552,438 | down |
|  |  | AT1G78580 | *TPS1* | 29,552,339 – 29,559,811 | down |
| Chr 1, Pos 29,605,216 | 9.39 x 10<sup>-9</sup> | AT1G78700 | *BEH4* | 29,599,348 – 29,601,682 | down |
|  |  | AT1G78730 | na | 29,607,688 – 29,609,441 | -- |
| Chr 4, Pos 9,567,764 | 1.33 X 10<sup>-8</sup> | AT4G16990 | *RLM3* | 9,558,570 – 9,565,555 | down |
|  |  | AT4G17000 | na | 9,566,876 – 9,569,630 | down |
| Chr 5, Pos 6,070,752 | 8.22 x 10<sup>-9</sup> | AT5G18320 | *PUB46* | 6,064,169 – 6,066,284 | -- |
|  |  | AT5G18340 | *PUB48* | 6,070,345 – 6,072,334 | up |
| Chr 5, Pos 6,080,536 | 3.18 x 10<sup>-9</sup> | AT5G18360 | *BAR1* | 6,079,767 – 6,083,077 | down |
|  |  | AT5G18370 | *DSC2* | 6,083,400 – 6,089,041 | down |

<sup>a</sup> Single-nucleotide polymorphisms (SNPs) are specified by their position on the respective chromosome according to https://www.1001genomes.org. <sup>b</sup> Genes within ± 5 kb of the detected SNPs that were expressed in 9-day-old Col-0 seedlings. <sup>c</sup> Heat-dependent transcriptional regulation in 9-day-old Col-0 seedlings following exposure to 37 °C for 1 hour as determined by RNA-seq. See Supplemental Table S2 for expression data. na: not available

Among these genes, *PLANT U-BOX 48* (*PUB48*, AT5G18340) was associated with the most significant SNP on chromosome 5 (position 6,070,752, *p* = 8.22 x 10^-9^). *PUB48* encodes an E3 ubiquitin ligase that has been linked to abiotic stress responses in Arabidopsis and whose transcription is induced by high temperature (Adler *et al*., 2017, Peng *et al*., 2019). Moreover, *pub48* mutant seeds are highly sensitive to heat suggesting a positive role for PUB48 during plant thermotolerance (Adler *et al*., 2017, Peng *et al*., 2019). Since we were most interested in the identification of genes that had not been linked to heat stress responses previously, we did not characterise *PUB48* in more detail.

The most significant SNP on chromosome 1 (position 29,547,498, *p* = 3.51 x 10^-9^) was located in close proximity to two genes that are directly related to carbohydrate metabolism, i.e., *RHAMNOSE BIOSYNTHESIS 1* (*RHM1*, AT1G78570) and *TREHALOSE-6-PHOSPHATE SYNTHASE 1* (*TPS1*, AT1G78580). RHM1 catalyses the conversion of UDP-glucose to UDP-rhamnose and provides the substrate for both flavonol glycosylation as well as the synthesis of cell wall polysaccharides (Ringli *et al*., 2008, Saffer and Irish, 2018). TPS1, on the other hand, catalyses the formation of trehalose 6-phosphate (T6P) from UDP-glucose and glucose 6-phosphate (Fichtner and Lunn, 2021). T6P serves as a homeostatic regulator of sucrose production and an indicator of endogenous sucrose availability thereby synchronizing plant growth and energy status *in planta* (Lunn *et al*., 2006, Fichtner and Lunn, 2021). Moreover, *TPS1* has been linked to abiotic stress responses such as drought stress (Figueroa and Lunn, 2016) and its catalytic product T6P also appears to regulate thermomorphogenic growth responses under moderately elevated temperatures (Hwang *et al*., 2019).

Therefore, we examined the genomic region around this SNP (position 29,547,498 on chromosome 1) in more detail. Out of the 250 accessions 19 carried the minor allele (A) and 231 the major allele (G) in this position. Accessions with the minor allele contained on average twice as much raffinose after the high temperature treatment as the accessions carrying the major allele (Figure 1c). This difference was less pronounced, yet also statistically significant, at the control temperature. However, we did not observe any association between raffinose levels at 22 °C and this genomic region on chromosome 1 across the 250 accessions in our mapping population (Figure S6).

Although we had observed a positive correlation between raffinose concentrations at 32 °C and latitude of origin, no clear pattern was apparent from the geographic distribution of the two alleles at this genomic position (Figure S7). Subsequent analysis of the linkage disequilibrium (LD) showed that several SNPs inside both *RHM1* and *TPS1* were in strong LD with the lead SNP (Figure 1b). Moreover, the transcription of both genes is significantly downregulated in response to heat (Figure S8). Based on these observations, *RHM1* and *TPS1* were selected as candidate genes for further analyses.

### *TPS1* affects the biosynthesis of raffinose family oligosaccharides

To establish a functional role of *RHM1* and/or *TPS1* for warm temperature-dependent raffinose accumulation, soluble carbohydrates were analysed in the *rhm1-2* missense mutant encoding an RHM1 isoform that does not show any enzymatic activity *in vitro* (Diet *et al*., 2006), and two TILLING mutants containing weak alleles of *tps1, tps1-11* and *tps1-12* (Gómez *et al*., 2010), under our phenotyping conditions. While *rhm1-2* seedlings showed the expected increase in raffinose levels after the 32 °C treatment, no statistically significant differences were observed in comparison to wild-type seedlings at either temperature (Figure S9). By contrast, both *tps1-11* and *tps1-12* accumulated more than twice as much raffinose after exposure to elevated temperature compared to their respective wild type Landsberg *erecta* (L*er*) (Figure 2c). Both lines also contained significantly higher raffinose levels under control conditions. Since raffinose is an intermediate in the heat-responsive RFO biosynthetic pathway (Figure 2a), we also quantified the two RFO metabolites galactinol and stachyose in the *tps1* mutant lines. Both TILLING lines accumulated significantly higher amounts of the raffinose precursor galactinol, especially after the heat treatment. The *tps1-11* mutant also displayed higher concentrations of the raffinose-derived tetrasaccharide stachyose after the 32 °C treatment compared to the wild type (Figure 2b and d). These results suggested that metabolic flux through the RFO biosynthetic pathway was increased in the *tps1* TILLING lines.

**Figure 2.**
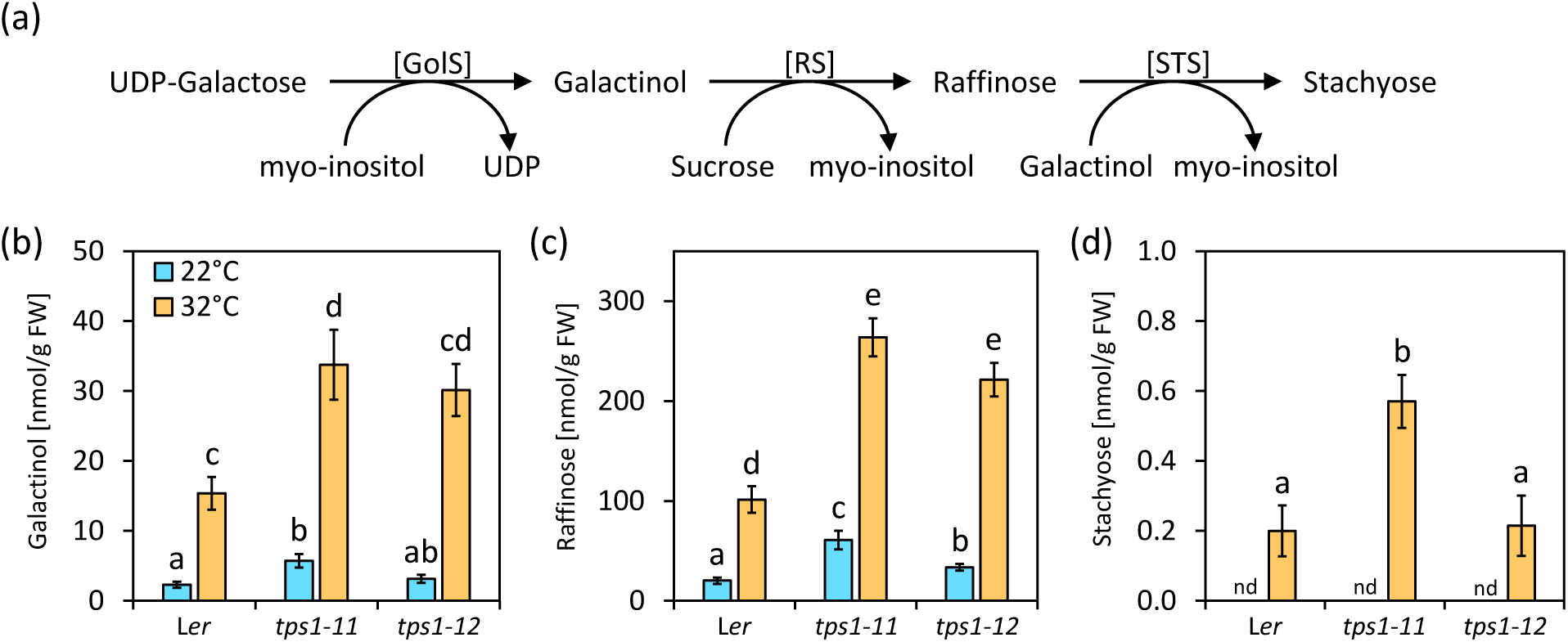
*TPS1* affects the biosynthesis of raffinose family oligosaccharides under moderate heat stress. (a) Schematic representation of the raffinose family oligosaccharide biosynthetic pathway. GolS: galactinol synthase; RS: raffinose synthase; STS: stachyose synthase. (b-d) Concentrations of galactinol, raffinose and stachyose in 14-day-old seedlings of Landsberg *erecta* (L*er*), *tps1-11* and *tps1-12* that were exposed to 32 °C for 4 hours (orange) or maintained at 22 °C as control (blue). Data are means ± SE, *N* = 6. Statistically significant differences are indicated by different letters above the bars (two-way ANOVA followed by a Tukey HSD test, *p* < 0.05). nd: not detected.

Since a previous study had reported higher levels of other soluble carbohydrates for the two TILLING lines (Gómez *et al*., 2010), we also quantified fructose, glucose and sucrose in *tps1-11* and *tps1-12* under our experimental conditions, and both TILLING lines tended to contain higher levels of these three sugars (Figure S10). Consequently, the calculated total amounts of soluble carbohydrates were also significantly higher in *tps1-11* and *tps1-12* compared to L*er* (Figure S10). In conclusion, our data support a role for *TPS1* not only in the regulation of carbohydrate metabolism under non-stress conditions, but also in the accumulation of warm temperature-dependent RFOs, with reduced TPS1 activity resulting in the over-accumulation of RFOs.

### Different TPS1 isoforms differentially affect carbohydrate metabolism during heat stress

Given that the *tps1* TILLING lines responded more strongly to elevated temperatures in terms of raffinose accumulation we next wanted to investigate whether *TPS1* is also involved in other temperature-dependent responses, in particular at more stressful temperatures. Although the two TILLING lines, *tps1-11* and *tps1-*12, had been backcrossed to L*er* (Gómez *et al*., 2010), we noticed that both lines still carried a significant fraction of Col-0-specific SNPs in their genomes. Based on RNA-sequencing (RNA-seq) reads we estimated the contribution of Col-0 to the *tps1-11* and *tps1-12* genetic backgrounds to be around 50% and 38%, respectively (Table S3).

Therefore, we decided to continue our functional analyses using two different *TPS1* complementation lines in the Col-0 *tps1-1* null mutant background that have been characterized recently (Fichtner *et al*., 2020). The line *TPS1[ΔN]* was generated by transforming *tps1-1* with a truncated *TPS1* gene encoding a version of the TPS1 protein that lacks the N-terminal domain. The N-terminal domain harbours a nuclear localisation signal and also possesses an auto-inhibitory function that lowers the enzymatic activity of TPS1. Consequently, *TPS1[ΔN]* seedlings display slightly increased T6P levels and appear to phenotypically resemble *TPS1* overexpression lines (Fichtner *et al*., 2020). By contrast, complementation of *tps1-1* with a construct lacking both the N-and C-terminal domains (denoted as *TPS1[ΔNΔC]*), resulted in plants that phenotypically resemble *tps1* knockdown mutants most likely due to a disruption of T6P signalling (Fichtner *et al*., 2020).

Given the involvement of TPS1 in the regulation of carbon partitioning (Fichtner and Lunn, 2021), we first assessed whether carbohydrate metabolism was altered in the two complementation lines during long-term heat stress. To this end, endogenous levels of transitory starch, sucrose and raffinose were quantified in 14-day-old seedlings after exposure to a high temperature treatment at 37 °C. Samples were taken after 24 and 39 hours into the heat treatment, corresponding to the end of the day and the end of the night, respectively (Figure 3a).

**Figure 3.**
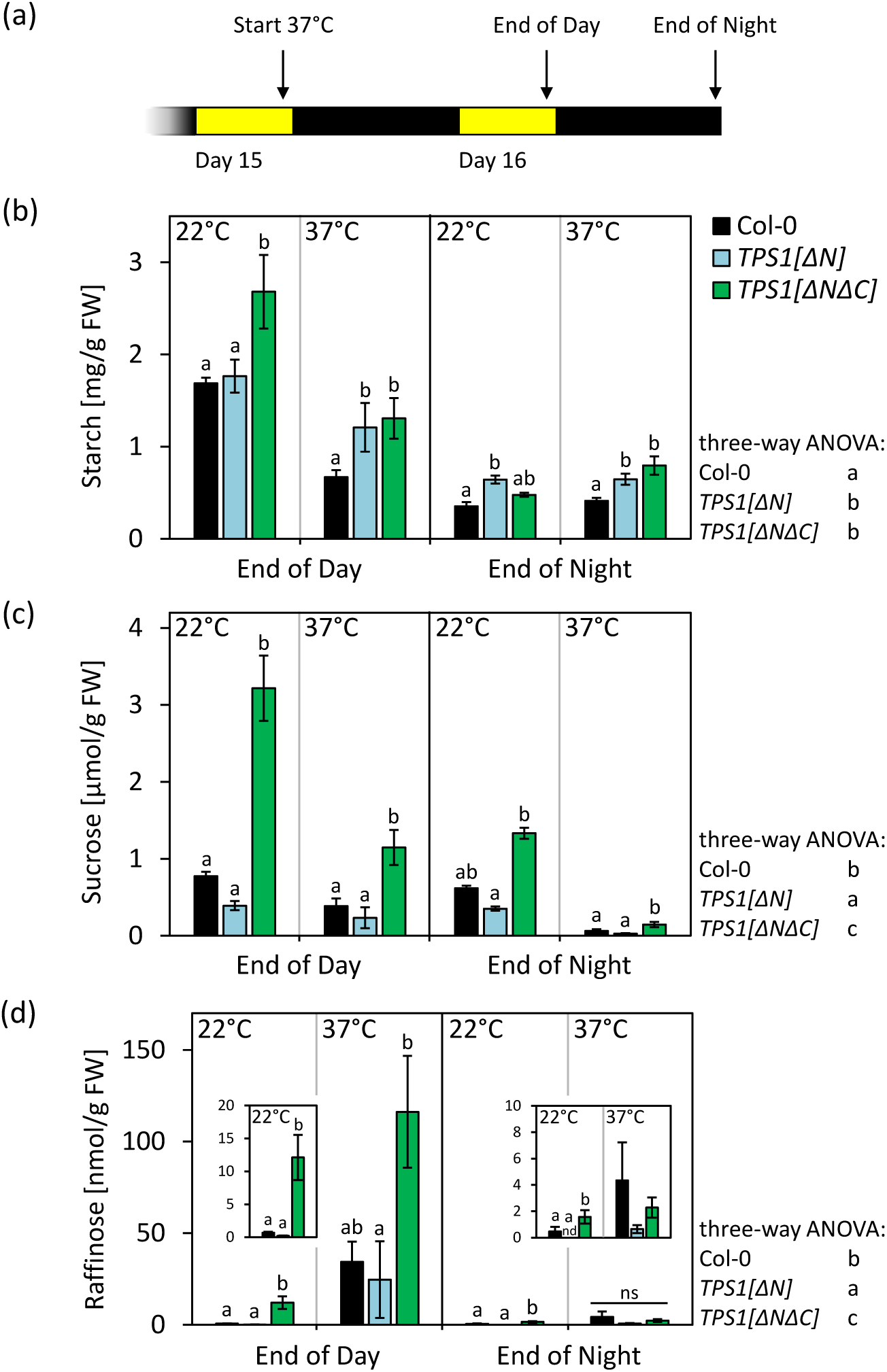
Complementation of *tps1-1* null mutants with different *TPS1* isoforms has contrasting effects on carbohydrate metabolism during long-term heat stress. (a) Schematic representation of the experimental conditions. The heat treatment was started at the end of day 15 and samples were taken at the end of the next day, i.e., after 24 hours, and at the end of the following night, i.e., after 39 hours. (b-d) Transitory starch, sucrose and raffinose concentrations in Columbia-0 (Col-0, black), *TPS1[ΔN]* (blue) and *TPS1[ΔNΔC]* (green). Data are means ± SE, *N* = 5-6. Letters above bars indicate statistically significant differences between the three genotypes as determined by a three-way ANOVA followed by a Tukey HSD test within the individual time and temperature conditions (*p* < 0.05). Overall significant differences between genotypes are indicated next to the plots (three-way ANOVA followed by a Tukey HSD test, *p* < 0.05). ns: not significant. nd: not detected.

As expected, transitory starch levels in wild-type seedlings were strongly reduced at the end of the night compared to the end of the light period under control conditions (Figure 3b). Notably, heat exposure did have a strong effect on endogenous starch levels during the day but not during the night (Figure 3b). Accordingly, heat-treated wild-type seedlings contained only 40% of the transitory starch levels that were detected in control seedlings at the end of the day, but were indistinguishable from non-treated seedlings at the end of the following night. Starch concentrations in *TPS1[ΔN]* were similar to wild-type levels at the end of the day under control conditions. However, at the end of the night *TPS1[ΔN]* seedlings contained significantly more starch than Col-0, and this was also the case for both time points during heat exposure. Similarly, *TPS1[ΔNΔC]* also contained significantly higher amounts of starch during the heat treatment compared to the wild type. However, in contrast to *TPS1[ΔN]*, *TPS1[ΔNΔC]* seedlings, already displayed higher starch levels than wild-type seedlings at the end of the day under control conditions. Overall, transitory starch levels were significantly higher in both complementation lines in comparison to Col-0 as indicated by a three-factorial ANOVA (Figure 3b).

Concerning endogenous sucrose, a different pattern was observed. *TPS1[ΔNΔC]* also displayed higher amounts of sucrose than the wild type irrespectively of time of the day and temperature treatment (Figure 3c). In contrast, *TPS1[ΔN]* contained lower levels than the wild-type throughout the whole experiment (Figure 3c). Although these differences were not statistically significant for the individual time points, the three-factorial ANOVA yielded a highly significant effect of genotype on endogenous sucrose levels, with *TPS1[ΔN]* displaying statistically significantly lower sucrose levels than the wild type and *TPS1[ΔNΔC]* displaying significantly higher levels, respectively (Figure 3c).

Since warm temperature-induced raffinose accumulation was our focal trait for the identification of *TPS1*, we also determined whether the different complementation lines were differentially affected in raffinose biosynthesis during heat stress. For all three genotypes the expected heat-induced increase in raffinose levels was observed (Figure 3d). Notably, *TPS1[ΔNΔC]* contained approximately three times as much raffinose as the wild type and *TPS1[ΔN]* after a 24-hour treatment at 37 °C. However, the difference between *TPS1[ΔNΔC]* and the wild-type seedlings was not statistically significant (*p* > 0.05). By contrast, raffinose levels in *TPS1[ΔN]* were similar to those in the wild type. It is also worth noting that under heat stress conditions raffinose levels declined from the end of the day until the end of the following night irrespectively of plant genotype, suggesting that raffinose is actively being metabolised during the night. The same was observed under control conditions, albeit to a lesser extent since raffinose levels were much lower under these conditions in general. Nevertheless, *TPS1[ΔNΔC]* already displayed significantly higher raffinose levels compared to the wild type at 22 °C both at the end of day and the end of the night, respectively. *TPS1[ΔNΔC]* also contained higher amounts of galactinol and stachyose than the wild type following the heat treatment (Figure S11).

Taken together, our data suggest that a fully functional TPS1 is required to keep endogenous starch and sucrose levels in balance under high temperature stress, in addition to its known function for carbon partitioning under non-stress conditions. Furthermore, the data from the complementation lines also corroborate our earlier findings that decreased TPS1 activity results in an over-accumulation of RFOs.

### Changes in TPS1 activity are associated with differences in thermotolerance

Finally, we determined whether *TPS1* also affects thermotolerance at the whole-plant level. To do so, 14-day-old seedlings were exposed to a 37 °C heat treatment for 48 hours. Following a recovery period of two weeks under non-stress conditions, survival of the seedlings was assessed based on their capacity to grow new leaves. While on average 38% of the wild-type seedlings did survive the heat treatment, *TPS1[ΔN]* displayed a significantly lower survival rate of 12% (Figure 4). By contrast, *TPS1[ΔNΔC]*, showed a significantly higher survival rate of 76% (Figure 4). Thus, our observations suggest that changes in TPS1 activity and the concomitant alterations in T6P-dependent signalling affect plant thermotolerance. Moreover, the observed differences in thermotolerance were preceded by differences in carbohydrate metabolism with enhanced tolerance being positively correlated with sucrose but not transitory starch levels during heat stress.

**Figure 4.**
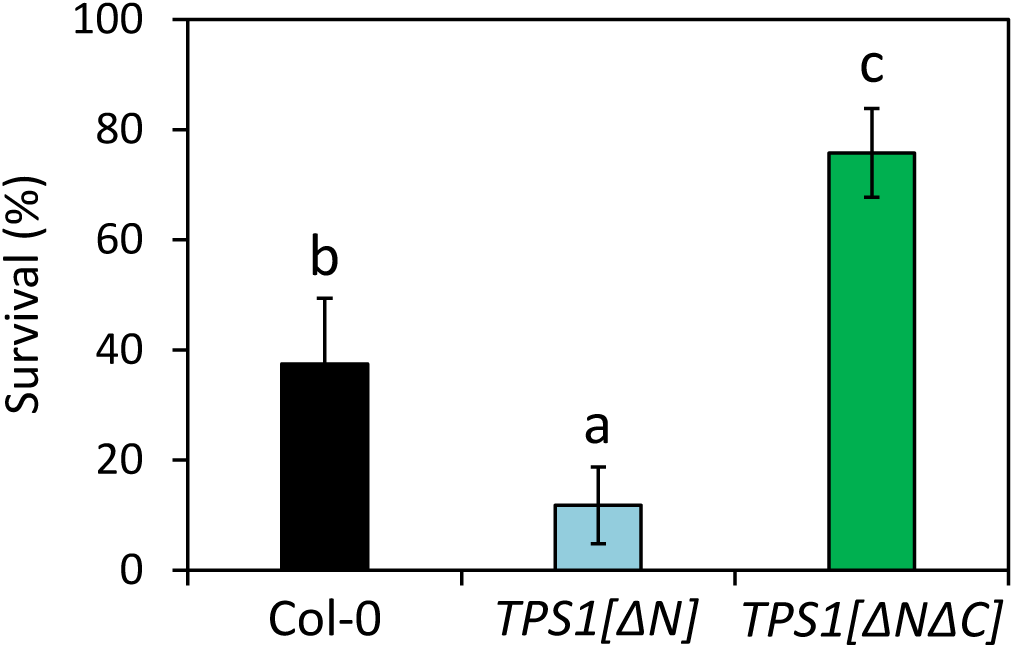
Complementation of *tps1-1* null mutants with different *TPS1* isoforms differentially affects long-term thermotolerance. Fourteen-day-old seedlings of Columbia-0 (Col-0, black), *TPS1[ΔN]* (blue) and *TPS1[ΔNΔC]* (green) were exposed to a 37 °C heat treatment for 48 hours. Following a 14-day recovery period under control conditions seedling survival was assessed based on the emergence of newly developed leaves. Data are means ± SE, *N* = 12. Different letters above bars indicate statistically significant differences (one way ANOVA followed by a Tukey HSD test, *p* < 0.05). Similar results were obtained in an independent experiment (Figure S12).

## Discussion

Surprisingly few studies have explored the intraspecific variation of responses towards high temperature stress with the aim to identify the underlying regulatory genes in Arabidopsis. In addition, most of these studies have focused on rather complex, macroscopic traits such as silique length, that might be the outcome of a similarly complex genetic architectures with many loci of small effect size contributing to the phenotype of interest (e.g. Bac-Molenaar *et al*., 2015, Thoen *et al*., 2017). However, such a large number of small-effect loci may impede the detection of statistically significant associations in GWAS (Korte and Farlow, 2013). Thus, the phenotyping of less complex traits, such as transcriptional or metabolic responses, with a presumably simpler genetic architecture, may facilitate the detection of significant associations and enhance the identification of genes involved in these responses.

In the present study, we have phenotyped 250 Arabidopsis accessions for their warm temperature-induced accumulation of the trisaccharide raffinose as a metabolic marker for heat stress resistance, based on its HSFA1-dependent regulation and its capacity to directly alleviate cellular damage evoked by heat (Panikulangara *et al*., 2004, Yan *et al*., 2022). In our phenotyping assay we observed a positive correlation between raffinose concentrations and latitude of origin, irrespectively of the temperature treatment, indicating that accessions from colder climates accumulate higher amounts of this metabolite. This notion was further supported by the observation that raffinose concentrations were negatively correlated, albeit weakly, with several temperature-related bioclimatic variables (Figure S4; Fick & Hijmans, 2017). While this might appear counterintuitive to the concept of raffinose serving as a metabolic marker for high temperature responses, this has also been observed for other heat stress-related traits such as HSP101 accumulation (Tonsor *et al*., 2008). One interpretation of these observations is that accessions from warmer climates with extended periods of high temperatures avoid excessive heat through early flowering rather than enhanced stress resistance, while accessions from cooler climates respond more strongly to occasional heat exposure (Wolfe and Tonsor, 2014). In addition, the latitudinal cline we observed fits with a recent study that investigated the natural variation of the primary metabolome in response to low temperature across 241 Arabidopsis accessions, in which a positive correlation between raffinose levels and latitude of origin was reported (Weiszmann *et al*., 2020). Unfortunately, the latter study did not report the genetic architecture underlying the observed variation.

In our genome-wide association study, phenotypic variation of warm temperature-induced raffinose accumulation was strongly associated with a region at the end of chromosome 1 that includes the *TPS1* locus. We subsequently provided several lines of evidence that *TPS1* is not only the causal gene underlying the observed variation in raffinose levels after a moderate temperature increase, but also affects metabolic responses and plant performance under more stressful temperature conditions. First, mutant lines with reduced TPS1 activity over-accumulated raffinose under our phenotyping conditions (Figure 2). Second, mutant lines with disrupted T6P signalling also accumulated higher amounts of transitory starch and several soluble carbohydrates during long-term exposure to 37 °C (Figure 3 and Figure S13). Third, impaired T6P signalling was associated with improved long-term thermotolerance, while higher TPS1 activity was accompanied by a reduction in thermotolerance (Figure 4 and Figure S14).

*TPS1* is expressed at different developmental stages and in various cell types such as guard cells and leaf vascular bundles as well as different cell types within the shoot apical meristem (Fichtner *et al*., 2020). Enzymatic TPS1 catalyses the formation of T6P from UDP-glucose and glucose 6-phosphate and is involved in embryo development, flowering and shoot branching (Fichtner and Lunn, 2021). Although noncatalytic functions of TPS1 have also been suggested, most of its functions have been attributed to the roles of its catalytic product T6P in sucrose signalling and homeostasis, which have been integrated in the so-called sucrose-T6P nexus model (Yadav *et al*., 2014). According to this model, T6P serves both as an indicator and regulator of endogenous sucrose levels *via* an intricate signalling network that regulates the partitioning of assimilated carbon during the day as well as the rate of starch degradation both during the day and during the night (Martins *et al*., 2013, Ishihara *et al*., 2022). It is proposed that T6P thereby mediates energy demand and supply between source and sink tissues (Figueroa and Lunn, 2016). In agreement with this, genetic perturbations of T6P synthesis or degradation strongly affect endogenous sucrose and starch concentrations (Martins *et al*., 2013, Figueroa *et al*., 2016, Fichtner *et al*., 2020).

Since *tps1* null mutants are non-viable due to an arrest during embryo development, we selected two *tps1* TILLING lines for our initial analyses, which were reported to contain strongly reduced T6P levels (Gómez *et al*., 2010). Both mutant lines over-accumulated several soluble carbohydrates during long-term heat stress and were able to maintain transitory starch levels that were similar to unstressed seedlings (Figure S13). By contrast, wild-type seedlings showed a strong reduction in transitory starch levels during heat exposure. These changes in carbohydrate metabolism were accompanied by an increase in long-term thermotolerance in both TILLING lines (Figure S14). Originally, these lines were generated through EMS-induced mutagenesis using seeds from a backcross between Col-0 and the *er105* fast-neutron-induced mutant and subsequently backcrossed three times with L*er* such that the residual contribution of Col-0 to their genetic backgrounds should be around 12.5 % (NASC, Till *et al*., 2003). However, in our RNA-seq analysis on their early transcriptional responses towards high temperature exposure we noticed that both TILLING lines contained a significantly higher proportion of Col-0-specific alleles than expected. Experiments with *tps1-11* and *tps1-12* should thus be interpreted with caution when using L*er* as wild-type control. Therefore, we sought additional evidence that *TPS1* is associated with high temperature stress in Arabidopsis.

Using *tps1-1* null mutant lines that were complemented with modified TPS1 isoforms, which have contrasting effects on T6P accumulation and signalling (Fichtner *et al*., 2020), most of our original observations using *tps1-11* and *tps1-12* were confirmed. While *TPS1[ΔN]* contains higher amounts of T6P and phenotypically resembles *TPS1* overexpressing lines, *TPS1[ΔNΔC]* phenotypically resembles *tps1* knockdown mutants in terms of the over-accumulation of certain carbohydrates but also their late flowering phenotype (Gómez *et al*., 2010, Fichtner *et al*., 2020). This has been attributed to the occurrence of two unidentified disaccharide-monophosphates in this line that may act as inhibitors of T6P signalling rather than changes in T6P levels themselves (Fichtner *et al*., 2020). Based on these observations, we hypothesised that *TPS1[ΔNΔC]* should also resemble the two *tps1* TILLING lines during high temperature stress.

Indeed, *TPS1[ΔNΔC]* also contained higher amounts of galactinol, raffinose and stachyose after the heat treatment compared to wild-type seedlings. The same was true for sucrose and transitory starch, suggesting that reduced TPS1 activity is indeed associated with higher accumulation of soluble carbohydrates and transitory starch during heat stress. In line with this, *TPS1[ΔN]* seedlings contained consistently lower levels of sucrose and raffinose although the differences compared to the wild type were less pronounced. Unexpectedly, however, heat stressed *TPS1[ΔN]* seedlings contained higher starch levels that were comparable to the ones detected in *TPS1[ΔNΔC]* during the heat treatment both at the end of the light period and the end of the night.

Transitory starch is synthesised during the day and degraded during the night to provide carbon and energy in the absence of light, and T6P affects both processes. As an indicator of sucrose status, it diverts the flux of photoassimilates towards the synthesis of organic and amino acids when sucrose levels are sufficiently high during the day (Figueroa *et al*., 2016). During the night, increasing T6P levels slow down the rate at which starch is being degraded (Martins *et al*., 2013, Dos Anjos *et al*., 2018). Thus, the enhanced transitory starch and sucrose levels in *TPS1[ΔNΔC]* seedlings at the end of the day under control conditions might be considered a direct outcome of impaired T6P signalling in these plants leading to a reduced flux of photoassimilates towards the synthesis of organic and amino acids and increased availability for the synthesis of carbohydrates. In the same manner, increased starch and sucrose levels in the heat stressed seedlings might also be the result of altered photoassimilate partitioning and a redirection of the assimilated carbon flux towards carbohydrate formation. However, based on the transcriptomic data for the TILLING lines we would like to suggest an alternative hypothesis.

In wild-type seedlings, high daytime temperatures lead to a decrease in the formation of transitory starch, and this has mainly been attributed to a decrease in carbon assimilation due to reduced photosynthetic capacity and increased photorespiration (Thalmann and Santelia, 2017). Thus, the enhanced accumulation of sucrose and starch in heat stressed *TPS1[ΔNΔC]* seedlings might also be the outcome of a higher maintenance of photosynthetic capacity during the stress treatment. In line with this, the most pronounced differences between both *tps1* TILLING lines and L*er* in our transcriptomic analysis were observed for transcripts related to photosynthetic and tetrapyrrole metabolic processes and among them we detected many genes belonging to the *LIGHT-HARVESTING CHLOROPHYLL a/b BINDING PROTEIN* (*LHC*) superfamily (Figure S15). Members of the LHC family are involved in the transfer of light energy to the photochemical reaction centres and it has been speculated that their abundance might be related to thermotolerance (Wang *et al*., 2017). While *LHC* transcripts were strongly downregulated in L*er* seedlings within one hour of heat exposure, their expression was significantly less affected in both *tps1* lines, potentially indicating a sustained replacement of heat damaged LHC proteins during heat stress in these lines. As mentioned above, however, data from these experiments should be interpreted with caution. Nevertheless, the suggested association between improved maintenance of photosynthetic capacity during heat stress and the increased accumulation of transitory starch in genotypes with reduced T6P signalling warrants further investigation. Notably, heat stressed *TPS1[ΔN]* seedlings also contained higher starch levels, albeit, in the absence of an increase in sucrose. Recently, it has been shown that transitory starch is also being degraded to some extent during the day with degradation rates increasing towards the end of the light period (Ishihara *et al*., 2022). As during the night, T6P also inhibits starch degradation in the light (Ishihara *et al*., 2022). Thus, the enhanced starch levels in *TPS1[ΔN]* during the heat treatment and the simultaneous reduction in endogenous sucrose may be indicative of reduced starch mobilisation rates due to the elevated T6P concentrations in this line. As a consequence, the contrasting thermotolerance phenotypes of *TPS1[ΔN]* and *TPS1[ΔNΔC]* correlated best with endogenous sucrose levels during the heat treatment.

Plant thermotolerance depends on the onset of the HSR, which is a costly response in terms of energy and carbon demand due to the massive reconfiguration of the proteome (Guihur *et al*., 2022). Likewise, increased thermotolerance is associated with higher endogenous levels of soluble carbohydrates across different species and developmental stages such as creeping bentgrass (*Agrostis stolonifera*) or tomato (*Solanum lycopersicum*) pollen (Röth *et al*., 2015). Moreover, Arabidopsis seedlings are more sensitive to sugar starvation at elevated temperatures and their thermotolerance can be enhanced *via* supplementation with sucrose (Figure S14, Hwang *et al*., 2019, Olas *et al*., 2021). It is thus likely that the enhanced heat resistance of *TPS1[ΔNΔC]* is the result of the combined increase in energy reserves, i.e., in the form of transitory starch, and energy availability, i.e., in the form of sucrose. In agreement with this, the reduced thermotolerance phenotype of *TPS1[ΔN]* can be explained by a reduced energy availability despite higher starch reserves as compared to the wild type. In addition, increased levels of soluble carbohydrates may also directly contribute to the enhanced resistance *via* the mitigation of oxidative damage caused by the heat treatment (Bolouri-Moghaddam *et al*., 2010).

The contrasting thermotolerance phenotypes we observed may, however, also be affected indirectly by the observed changes in carbohydrate accumulation *via* alterations in energy signalling. Plants employ an intricate signalling network in addition to the postulated sucrose-T6P nexus to monitor their sugar and energy status and adjust their growth and development accordingly. Central to this network are the two antagonistically operating kinases TARGET OF RAPAMYCIN (TOR) and SUCROSE NON-FERMENTING 1 RELATED-KINASE 1 (SnRK1), both of which have been linked to abiotic stress responses (Rodriguez *et al*., 2019). While TOR promotes plant growth and development when energy supply is sufficient, SnRK1 represses energy-consuming processes and induces catabolic metabolism under energy limiting conditions (Dröge-Laser and Weiste, 2018). TOR activity itself is positively regulated by high glucose and sucrose levels, although it is unclear whether the TOR complex senses these metabolites directly or indirectly (Rodriguez *et al*., 2019). Recently, it has been shown that TOR overexpressing lines are more thermotolerant and that this is the outcome of TOR-mediated epigenetic regulation of heat stress-associated genes (Sharma *et al*., 2022). In line with this, the improved thermotolerance of *TPS1[ΔNΔC]* seedlings might be a direct consequence of enhanced TOR activity due to the increased sucrose and glucose levels in the heat stressed seedlings. At the same time, these seedlings may perceive a contradicting signal of low energy status deriving from the constitutively low T6P:sucrose ratio that might result in the activation of SnRK1 (Baena-González and Lunn, 2020). However, surprisingly little is known about any involvement of SnRK1-dependent signalling in thermotolerance. Given the high metabolic costs that are associated with the onset of the HSR, it is tempting to speculate that this response is tightly coordinated with the plant’s energy status potentially allowing for a direct regulatory function of T6P in thermotolerance.

### Concluding remarks

In this study, we identified *TPS1* as a modulator of long-term thermotolerance, most likely through its regulatory function in carbon partitioning and sucrose homeostasis. However, more work is needed to understand how *TPS1* impinges exactly on plant thermotolerance. Future work should specifically address whether the observed changes in thermotolerance are mediated by T6P and investigate the role of potential downstream regulators. This might provide a means to uncouple the enhanced heat resistance from the penalties on plant growth associated with defects in *TPS1*, and result in the identification of promising targets for the breeding of climate-resilient crops.

## Acknowledgements

We thank John Lunn (Max Planck Institute of Molecular Plant Physiology, Potsdam-Golm, Germany) for discussions and comments on the manuscript, Pamela Korte and Theresa Damm for cultivation of plants, and Maria Lesch for excellent technical assistance with the fluorometric starch assay. Furthermore, we would like to acknowledge the contribution of undergraduate students that participated in the experimental work: Paula Hausmann, Priska Mair, Christoffer Lutsch and Jannik Bäsmann. This work was supported by the Deutsche Forschungsgemeinschaft (DFG, German Research Foundation) – project number 416992417.

