## Supporting Figures for "Natural variation of warm temperature-induced raffinose accumulation identifies *TREHALOSE-6-PHOSPHATE SYNTHASE 1* as a modulator of thermotolerance"

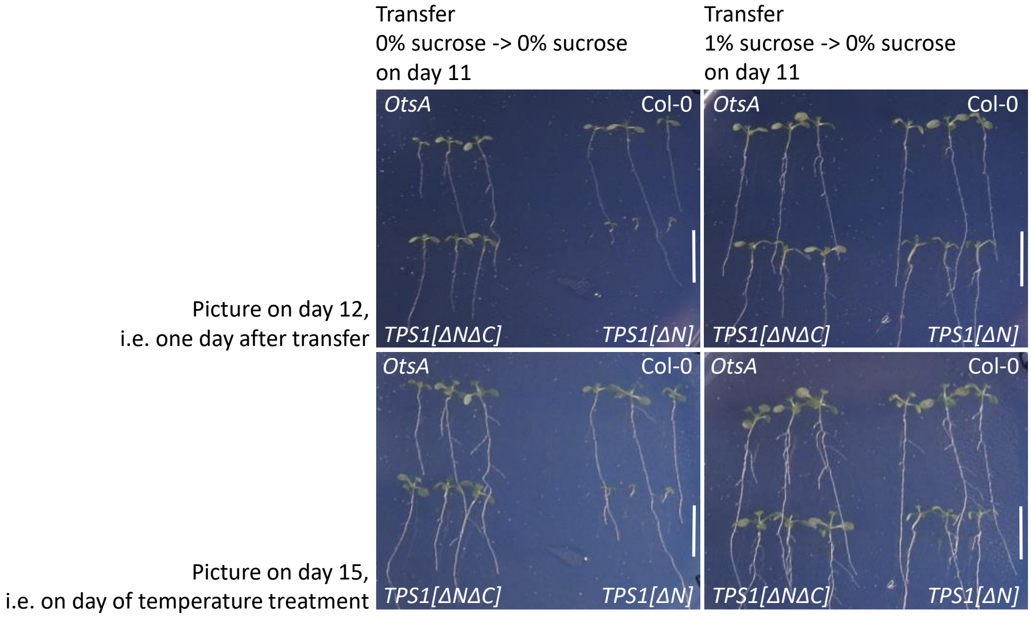


**Figure S1. Effect of sugar supplementation on *in vitro* growth of different *TPS1* complementation lines in the *tps1-1* null mutant background.** Seedlings of Columbia-0 (Col-0), *TPS1[ΔN]*, *TPS1[ΔNΔC]* and O*tsA* (Fichtner *et al.*, 2020) were grown under short day conditions (9:15 h light/dark; 22°C) on ½-strength MS-Agar, pH 5.7, either in presence or absence of 1% sucrose. After ten days all seedlings were transferred to sucrose-free ½-strength MS-Agar, pH 5.7, and grown under the same conditions for another four days. Pictures were taken on day 12 and day 15, i.e., one day after the transfer and on the day of the temperature treatments, respectively. Sucrose supplementation could rescue the otherwise poor growth of *TPS1[ΔN]*. Scale bars represents 1 cm.

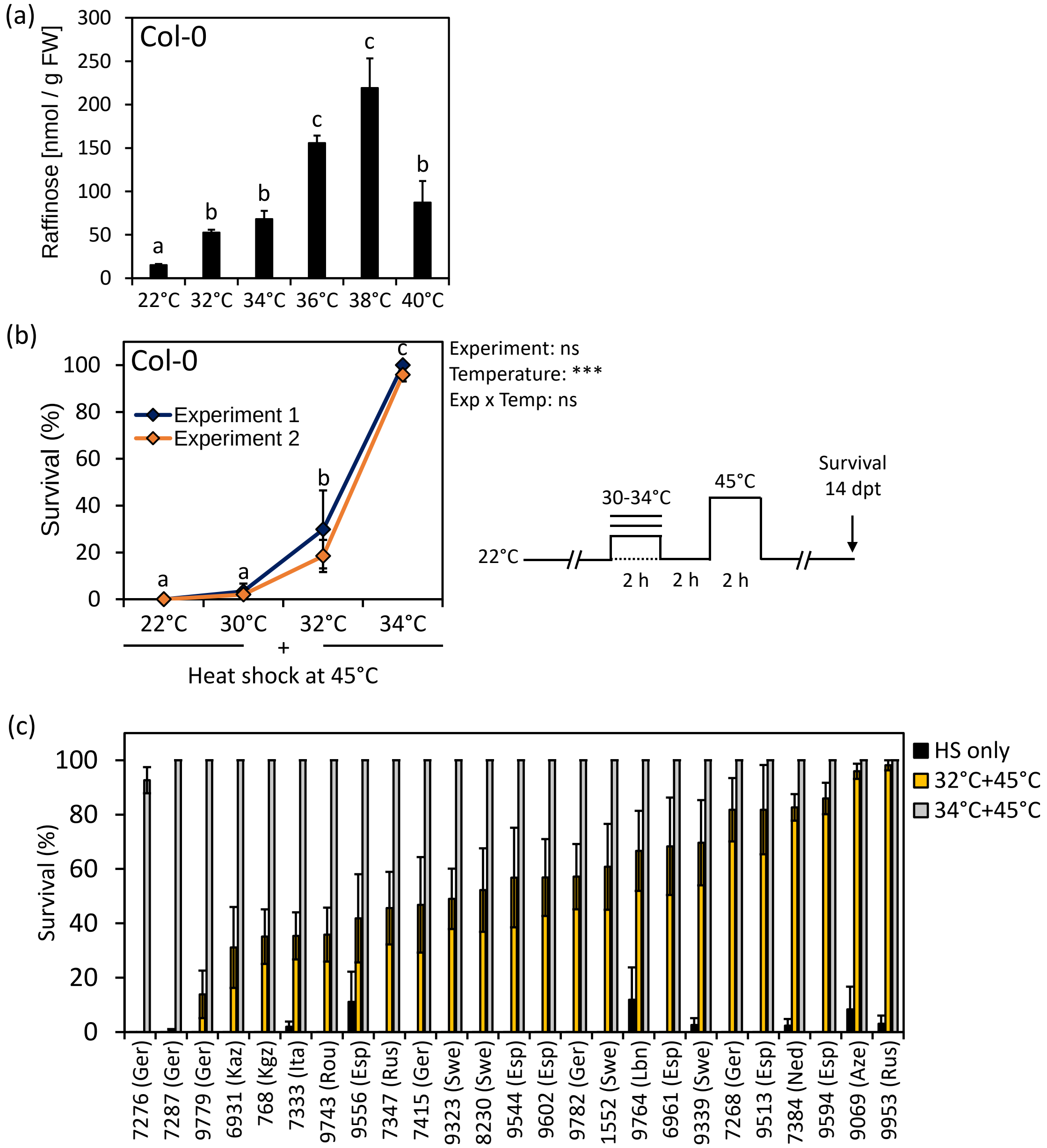


**Figure S2. Temperature-dependent effects on raffinose accumulation and short-term acquired thermotolerance.** (a) Effect of elevated temperatures on raffinose accumulation. Fourteen-day-old seedlings of Columbia-0 (Col-0) were exposed to various temperatures for 2 hours or maintained at 22 °C as control. Data are means ± SE. *N* = 4. Letters above bars indicate statistically significant differences (one-way ANOVA followed by a Tukey HSD test, $p < 0.05$). (b) Short-term acquired thermotolerance as a function of acclimation temperature. Fourteen-day-old seedlings of Col-0 were acclimated at different temperatures for 2 hours followed by a 2-hour lag phase at 22 °C prior to the final heat shock treatment at 45 °C. Seedling survival was determined after a 14-day recovery period under control conditions based on the emergence of newly developed leaves. Shown are means ± SE for two independent experiments. *N* = 5. Letters indicate statistically significant differences (two-way ANOVA followed by a Tukey HSD test, $p < 0.05$). Overall effects of the two factors 'Experiment' and 'Acclimation Temperature' as well as the interaction term ('Exp x Time') are given next to the plot (*** $p < 0.001$, ns: not significant). A schematic representation of the temperature protocol used is depicted on the right. (c) Natural variation of short-term acquired thermotolerance across 25 accessions of *Arabidopsis thaliana*. Fourteen-day-old seedlings of the various accessions were acclimated for two hours at 32 °C or 34 °C or not acclimated at all ('HS only') prior to the final heat shock at 45 °C. Seedling survival was determined after a 14-day recovery period under control conditions based on the emergence of newly developed leaves. Shown are means ± SE. *N* = 6-12 for acclimated seedlings, *N* = 3-6 for non-acclimated seedlings.

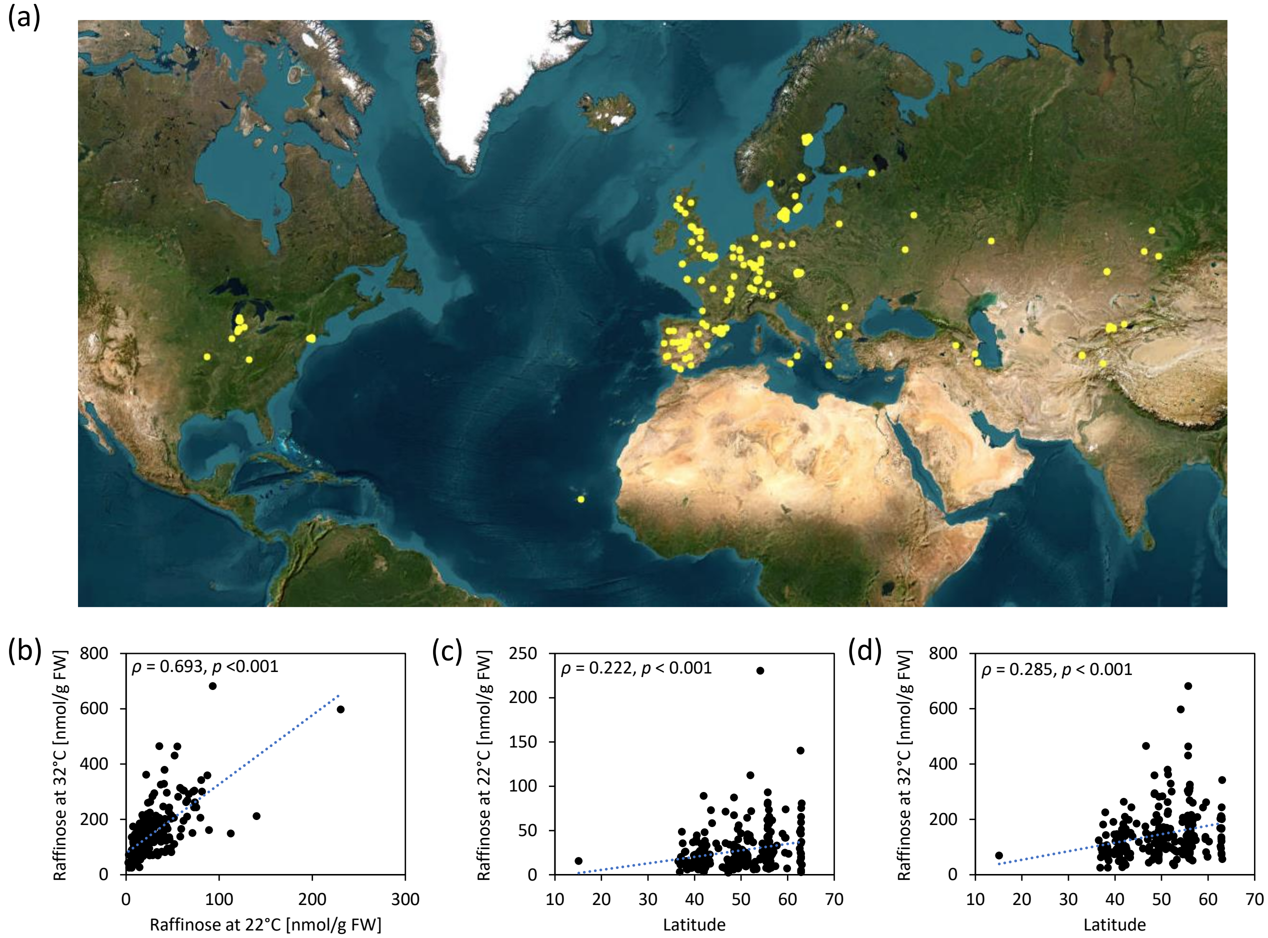


**Figure S3. Correlation between latitude of origin and raffinose accumulation across 250 *Arabidopsis thaliana* accessions.** (a) Geographic distribution of the 250 accessions used for the phenotyping. (b-d) Correlation between raffinose concentrations at 22 °C and after a 4-hour treatment at 32 °C, raffinose concentrations at 22 °C and latitude of origin, and raffinose concentrations at 32 °C and latitude of origin. Correlation analyses were done using Spearman's rank correlation coefficient *rho*.

a)

| | BIO1 | BIO2 | BIO3 | BIO4 | BIO5 | BIO6 | BIO7 | BIO8 | BIO9 | BIO10 | BIO11 |
|---|---|---|---|---|---|---|---|---|---|---|---|
| Raffinose at 22 °C | **-0.160** | -0.069 | **-0.132** | 0.044 | **-0.125** | -0.081 | 0.014 | -0.017 | -0.089 | **-0.126** | -0.094 |
| Raffinose at 32 °C | -0.102 | **-0.191** | -0.090 | -0.101 | **-0.196** | 0.025 | **-0.146** | 0.003 | -0.007 | **-0.151** | 0.006 |

b) BIO1: Annual Mean Temperature
BIO2: Mean Diurnal Range (Mean of monthly (max temp - min temp))
BIO3: Isothermality (BIO2/BIO7)
BIO4: Temperature Seasonality (standard deviation x 100)
BIO5: Max Temperature of Warmest Month
BIO6: Min Temperature of Coldest Month
BIO7: Temperature Annual Range (BIO5 - BIO6)
BIO8: Mean Temperature of Wettest Quarter
BIO9: Mean Temperature of Driest Quarter
BIO10: Mean Temperature of Warmest Quarter
BIO11: Mean Temperature of Coldest Quarter

**Figure S4. Correlation between raffinose concentrations at different temperatures and temperature-related bioclimatic variables across 250 *Arabidopsis thaliana* accessions.** (a) Spearman's rank correlation coefficients for eleven bioclimatic variables (Fick & Hijmans, 2017) and raffinose concentrations at 22 °C and 32 °C, respectively. Statistically significant correlations are highlighted in bold typeface. Bioclim data used were at a spatial resolution of 2.5 minutes. (b) Brief description of the 11 temperature-related bioclimatic variables.

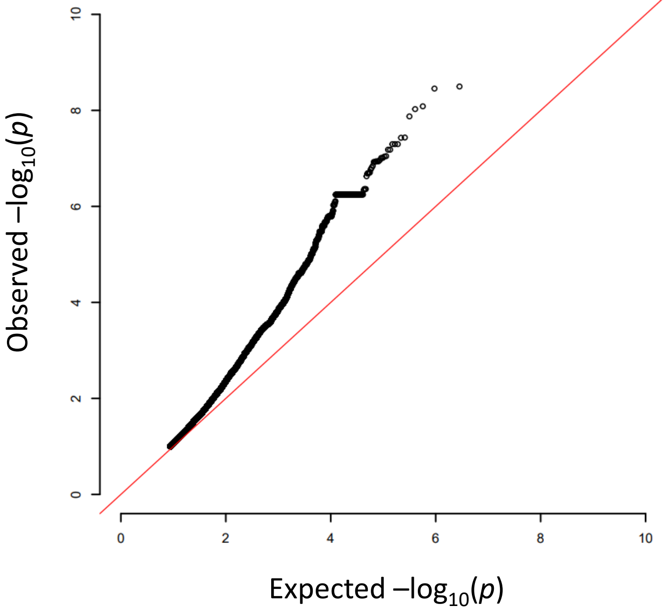


**Figure S5. QQ-plot for the genome-wide association study on raffinose concentrations after exposure to moderate heat at 32 °C.**

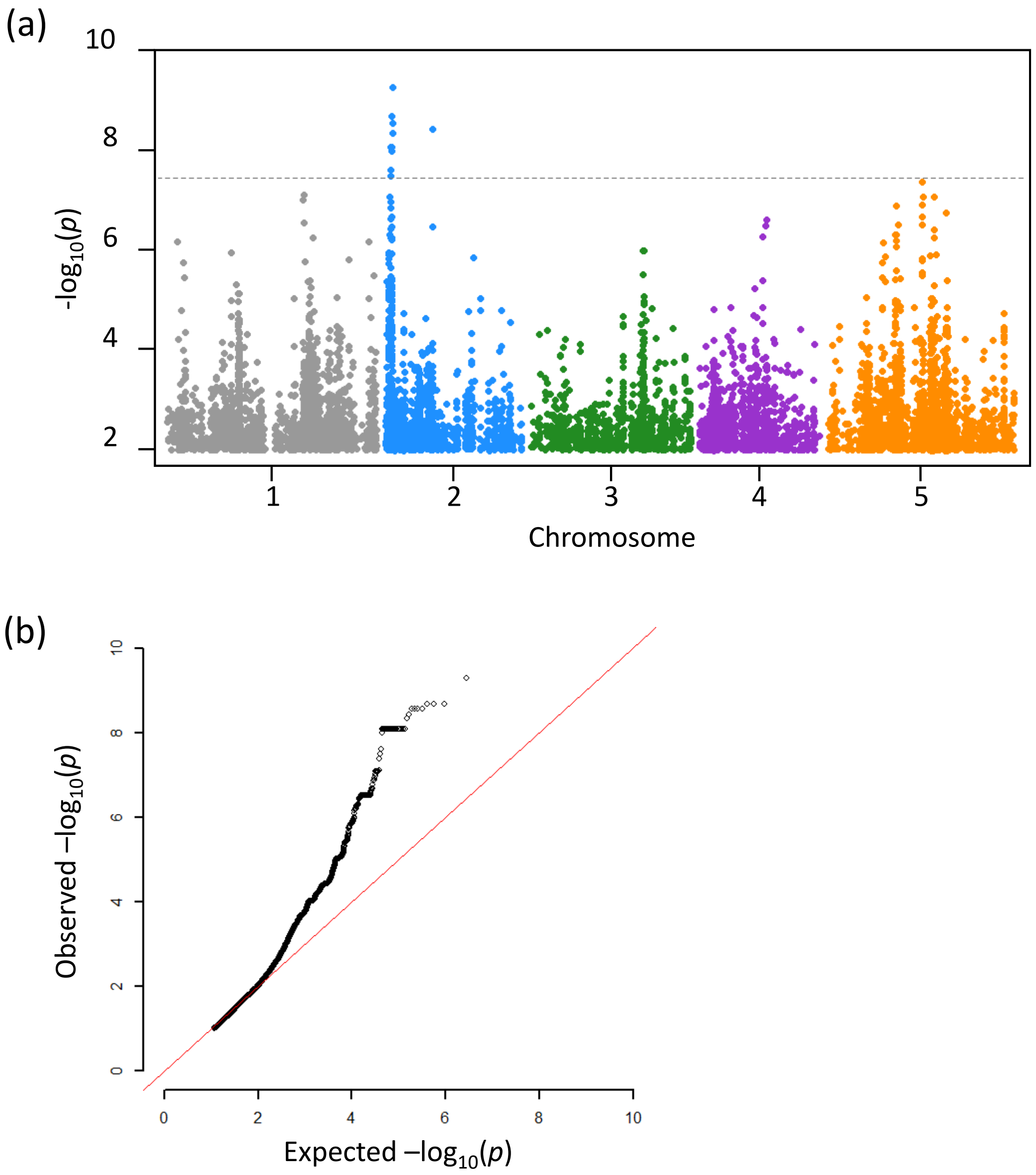


**Figure S6. Genome-wide association study on raffinose concentrations under control conditions.** (a) Manhattan plot depicting the statistical association between individual single-nucleotide polymorphisms (SNPs) and the variation in raffinose levels at 22 °C on a -$\log_{10}(p)$ scale. The dashed line indicates the Bonferroni-corrected significance threshold. (b) QQ-plot showing the relationship between the expected distribution of *p*-values and the observed *p*-value distribution of the genome-wide association study on raffinose concentrations at 22 °C.

(a)

(b)

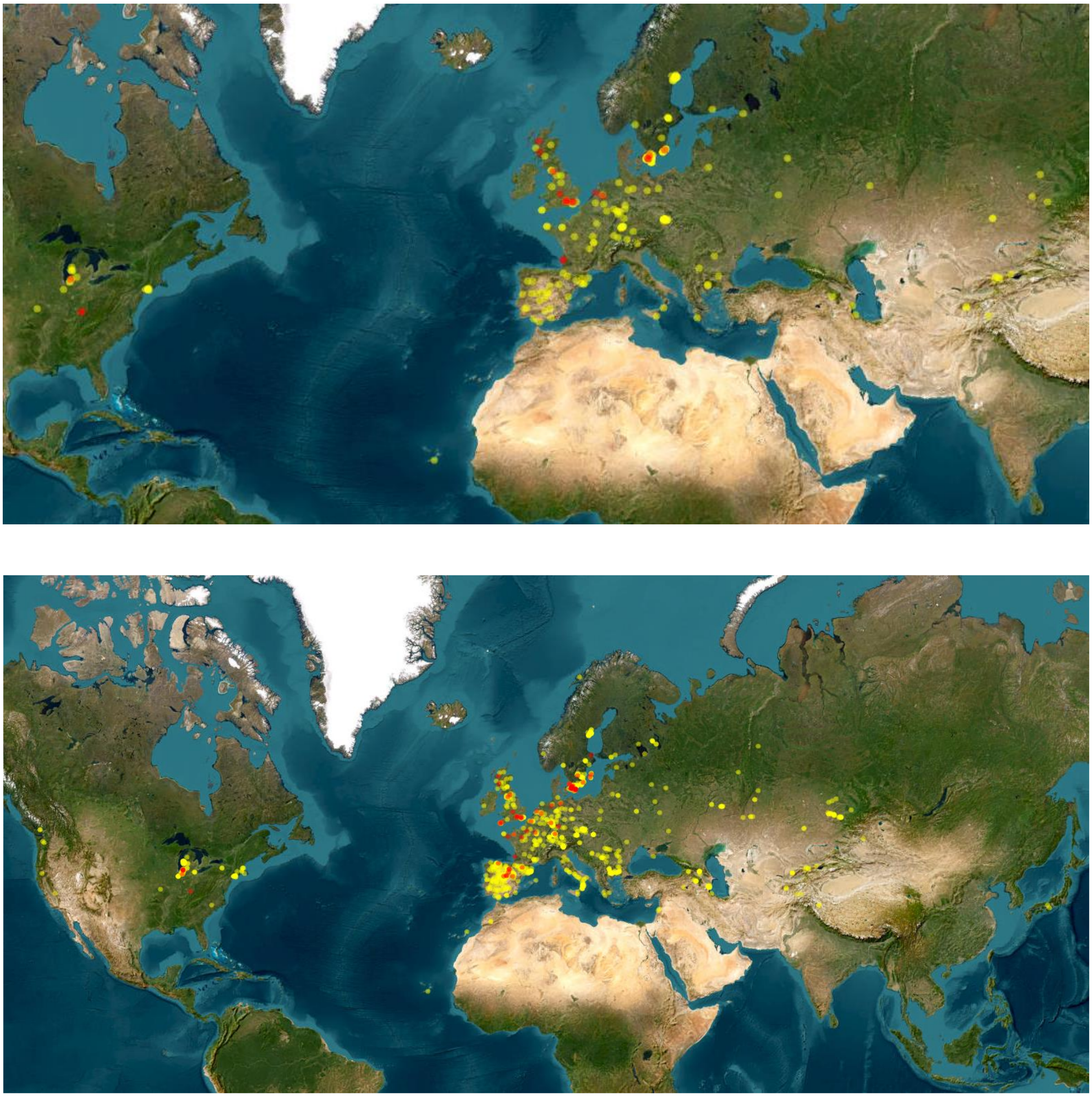

**Figure S7. Geographic distribution of the different alleles for the biallelic single-nucleotide polymorphism (SNP) at position 29,547,498 on chromosome 1.** (a) Geographic distribution of the two distinct alleles at position 29,547,498 on chromosome 1 within the mapping population of 250 accessions. (b) Geographic distribution of the distinct alleles across the 1135 accessions of the 1001 Genomes Project. The major allele (guanine) is depicted in yellow while the minor allele (adenine) is depicted in red.

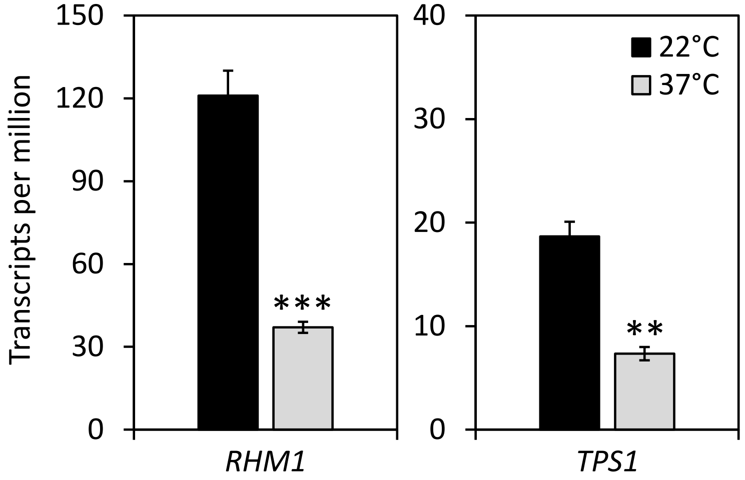


**Figure S8. Temperature-dependent expression of *RHAMNOSE BIOSYNTHESIS 1* (*RHM1*) and *TREHALOSE-6-PHOSPHATE SYNTHASE 1* (*TPS1*).** Expression data was derived from an unpublished in-house data set. In brief, Columbia-0 (Col-0) seedlings were grown on ½-strength MS-Agar, pH 5.7, for twelve days under long-day conditions (16:8 h, light/dark, 22 °C) before they were subjected to a 37 °C heat treatment for one hour or maintained at 22 °C as control. Library preparation and sequencing were performed by BGI Genomics, Shenzhen, China. Data are given in transcripts per million and shown as means ± SE, $N = 3$. Asterisks denote statistically significant differences compared to the control treatment (** $p < 0.01$, *** $p < 0.001$, Student's *t*-test).

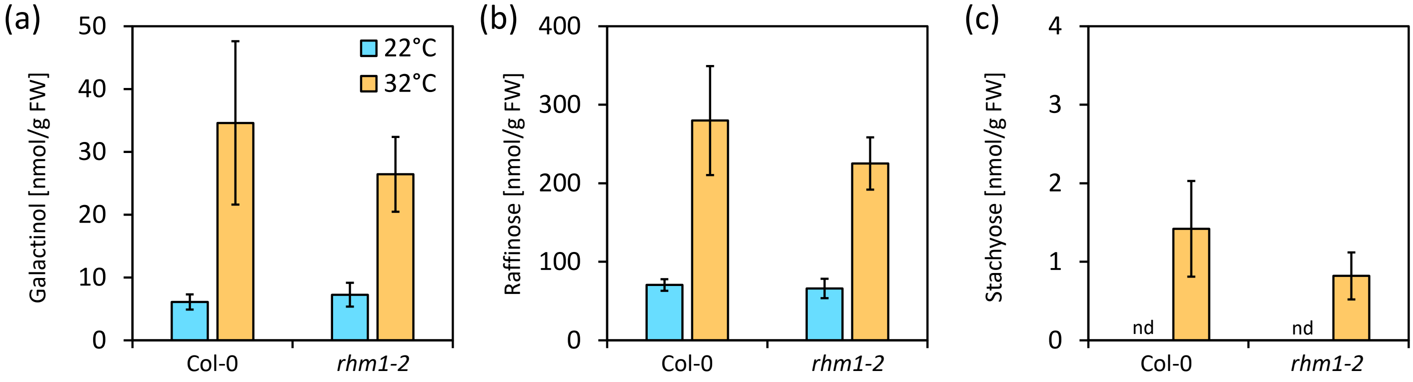


**Figure S9. Temperature-dependent accumulation of raffinose family oligosaccharides in *rhm1-2*.** Concentrations of galactinol (a), raffinose (b) and stachyose (c) in 14-day-old seedlings of Columbia-0 (Col-0) and *rhm1-2* that were exposed to 32 °C for 4 hours (orange) or maintained at 22 °C as control (blue). Data are means ± SE, *N* = 3 for Col-0 and *N* = 5 for *rhm1-2*. No statistically significant differences between *rhm1-2* and wildtype seedlings were detected. nd: not detected.

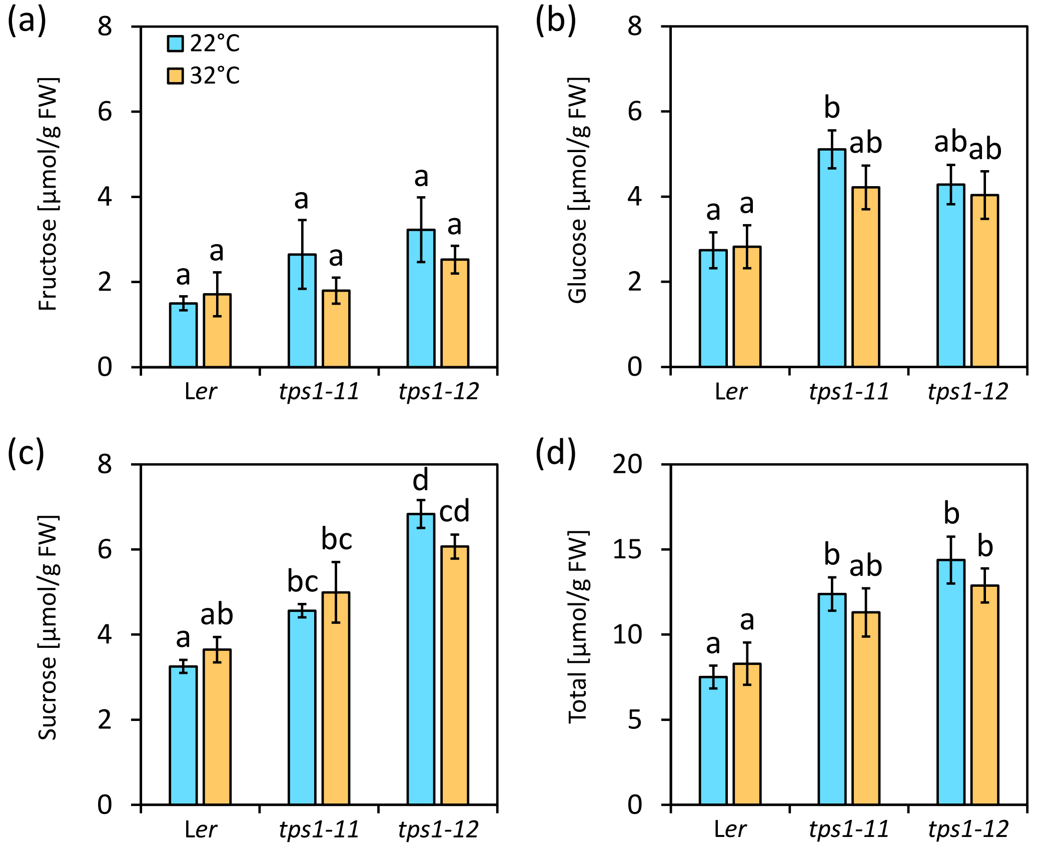


**Figure S10. Effect of short-term heat on soluble carbohydrates in *tps1-11* and *tps1-12*.** Concentration of fructose (a), glucose (b), sucrose (c) and total detected soluble carbohydrates (d) in 14-day-old seedlings of Landsberg *erecta* (L*er*), *tps1-11* and *tps1-12* after exposure to a moderately elevated temperature of 32 °C for 4 hours (orange). Control seedlings were maintained at 22 °C (blue). Data are means ± SE, $N = 6$. Statistically significant differences are indicated by different letters above bars (two-way ANOVA followed by a Tukey HSD test, $p < 0.05$). Total soluble carbohydrates is the sum of fructose, glucose, sucrose, galactinol, raffinose and stachyose.

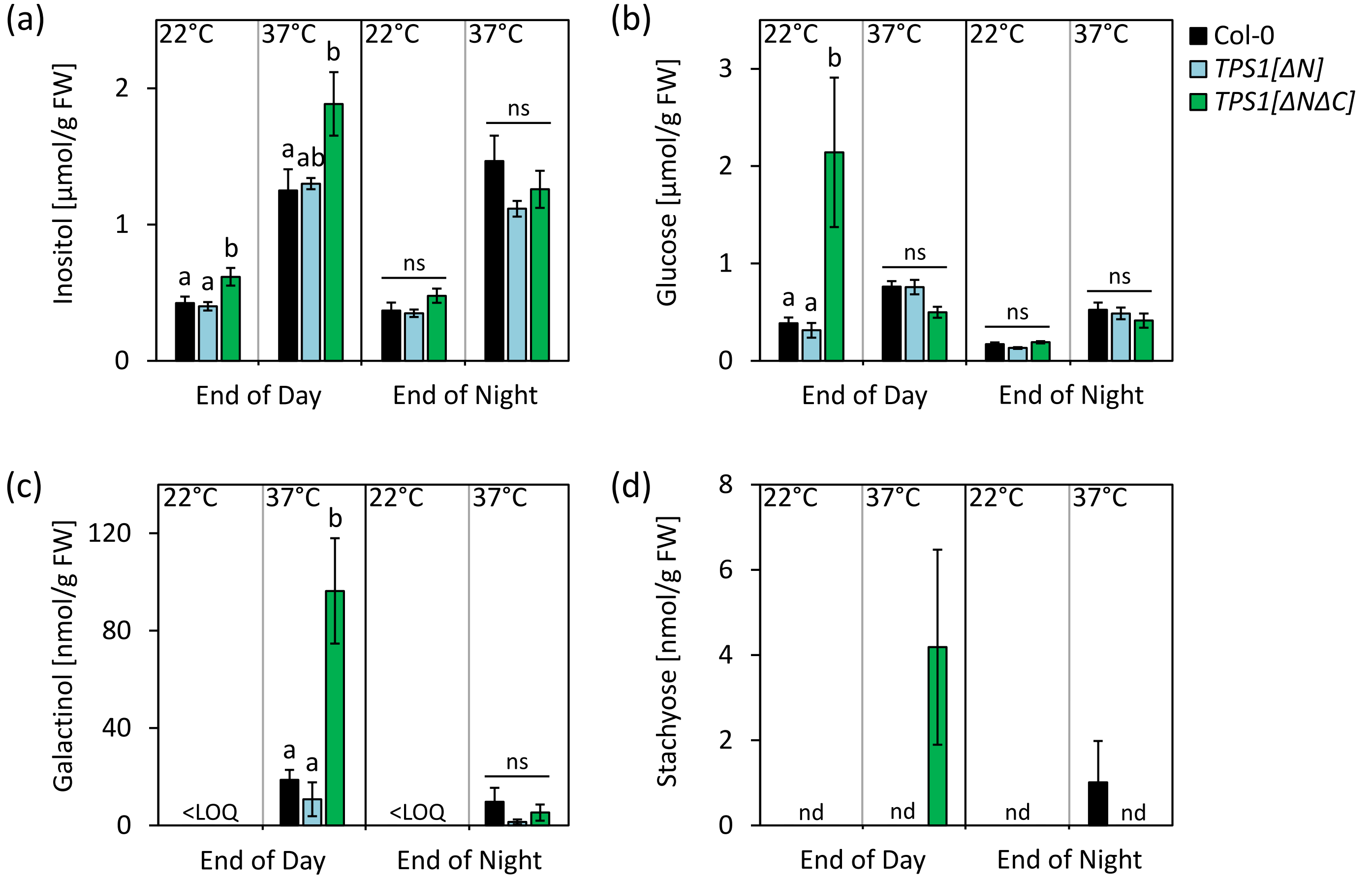


**Figure S11. Soluble carbohydrates in different *TPS1* complementation during long-term heat stress.** Concentrations of inositol (a), glucose (b), galactinol (c) and stachyose (d) in seedlings of Columbia-0 (Col-0, black), *TPS1[ΔN]* (blue) and *TPS1[ΔNΔC]* (green) that were exposed to 37 °C or maintained at 22°C as control. The 37 °C heat treatment was started at the end of day 15 (ZT 8) and samples were taken at the end of the next day, i.e., after 24 hours (ZT 8), and at the end of the following night, i.e., after 39 hours (ZT 23). Data are means ± SE, *N* = 5-6. Letters above the bars indicate statistically significant differences between the three genotypes within the individual time and temperature conditions as determined by a three-way ANOVA followed by a Tukey HSD test ($p < 0.05$). Note that for stachyose the statistical significance was not tested. nd: not detected; LOQ: limit of quantification; ns: not significant.

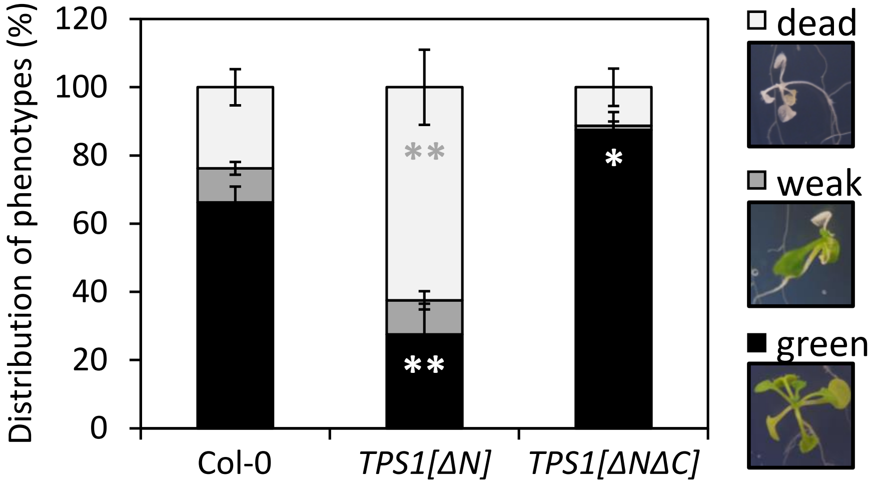


**Figure S12. Effect of long-term heat on seedling performance in different *TPS1* complementation lines.** (a) Long-term thermotolerance of Columbia-0 (Col-0), *TPS1[ΔN]* and *TPS1[ΔNΔC]*. Fourteen-day-old seedlings were exposed to a 37 °C long-term heat treatment for 72 hours. After a recovery period of 14 days under control conditions, seedlings were categorised as 'dead', 'weak' or 'green' according to the pictures shown on the right. The proportional distribution of the three categories was calculated within individual agar plates and is given as mean ± SE, $N = 8$. Asterisks indicate statistically significant differences (* $p < 0.05$, ** $p < 0.01$, paired $t$-test).

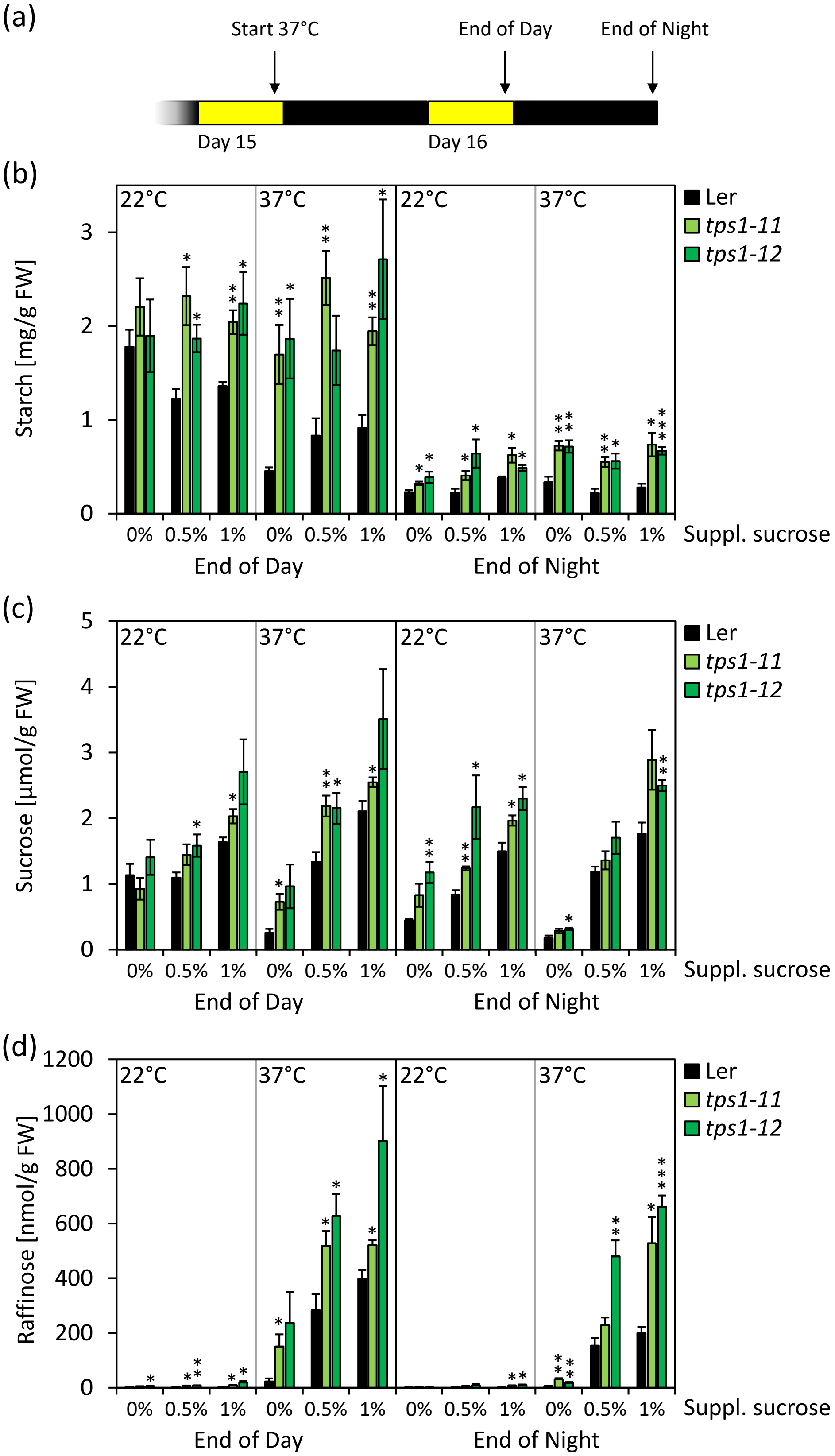


**Figure S13. Effect of sucrose supplementation on endogenous carbohydrates during long-term heat stress in *tps1-11* and *tps1-12*.** (a) Schematic representation of the experimental conditions. The 37 °C heat treatment was started at the end of day 15 (ZT 8) and samples were taken at the end of the next day, i.e., after 24 hours (ZT 8), and at the end of the following night, i.e., after 39 hours (ZT 23). (b-d) Transitory starch, sucrose and raffinose concentrations in seedlings of Landsberg *erecta* (L*er*, black), *tps1-11* (light green) and *tps1-12* (dark green) that were exposed to 37 °C or maintained at 22°C as control. Data are means ± SE, $N = 4$. Asterisks indicate statistically significant differences compared to the wildtype (* $p < 0.05$, ** $p < 0.01$, *** $p < 0.001$, Student's $t$-test). All seedlings were grown on ½-strength MS-Agar containing 1% sucrose for ten days before they were transferred to ½-strength MS-Agar plates supplemented with the indicated sucrose concentrations.

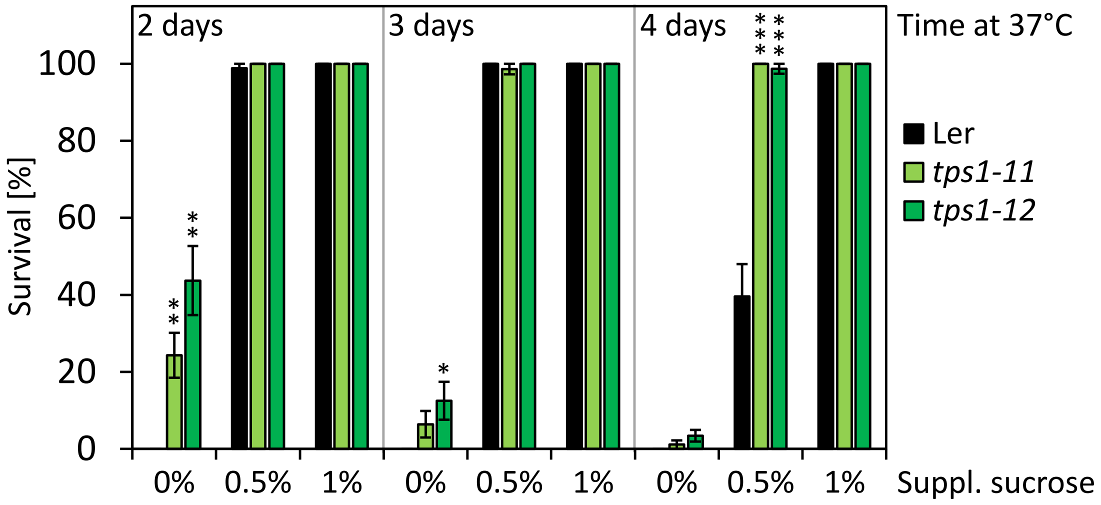


**Figure S14. Effect of sucrose supplementation on long-term thermotolerance of *tps1-11* and *tps1-12*.** Survival of 14-day-old seedlings of Landsberg *erecta* (Ler), *tps1*-11 and *tps1*-12 that were exposed to a 37 °C heat treatment for two, three and four days, respectively. Following a fourteen-day recovery period under control conditions seedling survival was assessed based on the emergence of newly developed leaves. All seedlings were grown on ½-strength MS-Agar containing 1% sucrose for ten days before they were transferred to ½-strength MS-Agar plates supplemented with the indicated sucrose concentrations. Data are means ± SE, $N$ = 6-7. Asterisks indicate statistically significant differences compared to wildtype seedlings (* $p < 0.05$, ** $p < 0.01$, *** $p < 0.001$, paired $t$-test).

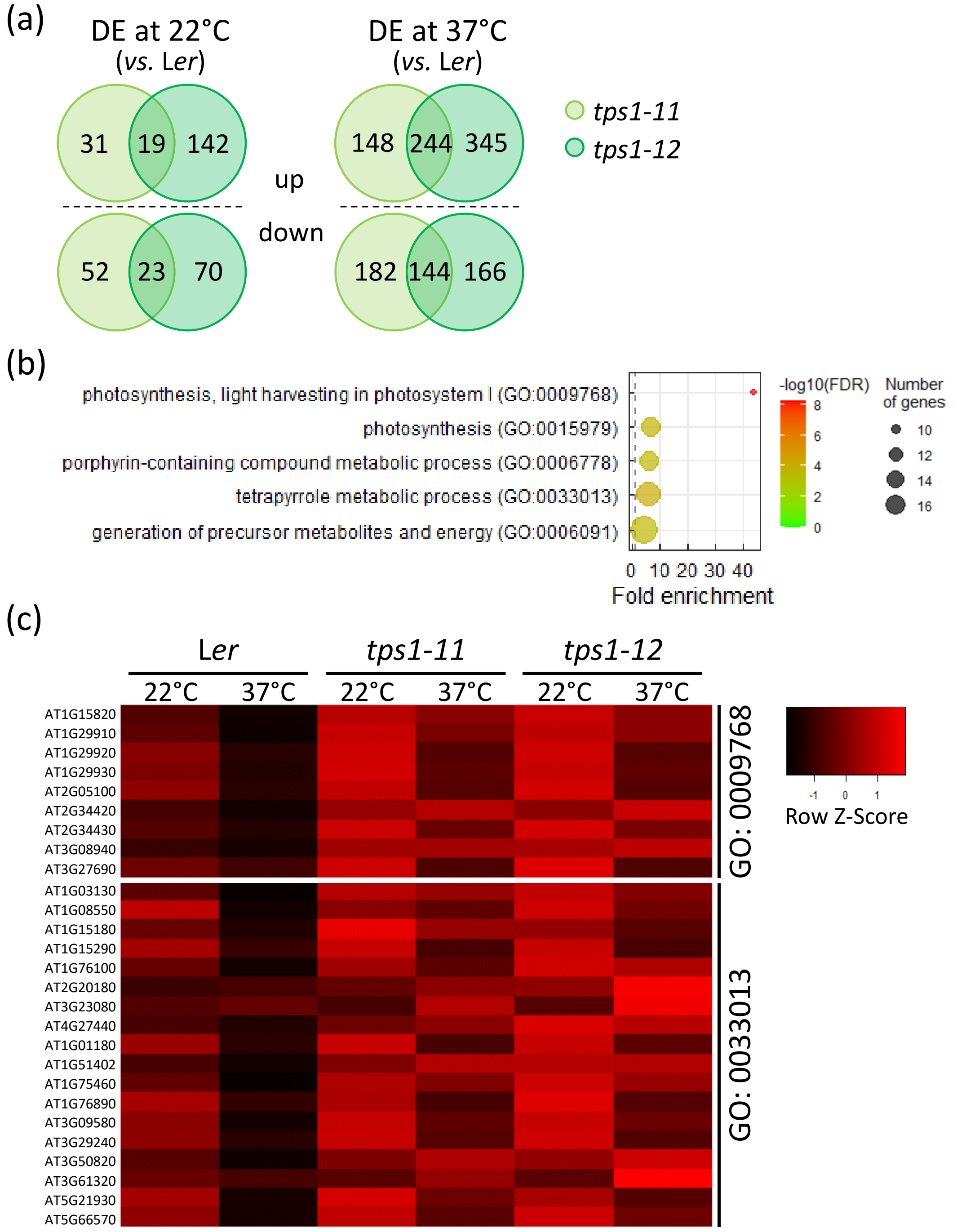


**Figure S15. Differences in the early transcriptional response of *tps1-11* and *tps1-12* towards heat stress.** (a) Venn diagrams showing the numbers of up- and down-regulated genes in 14-day-old seedlings of *tps1-11* and *tps1-12* compared to Landsberg *erecta* (L*er*) after heat treatment at 37 °C for one hour and under control conditions (22 °C), respectively. (b) Gene Ontology (GO) term enrichment analysis for the 244 genes that were more highly expressed in both *tps1-11* and *tps1-12* after the heat treatment. (c) Expression profiles of differentially expressed genes within the two GO terms 'photosynthesis, light harvesting in photosystem I' (GO:0009768) and 'tetrapyrrole metabolic process' (GO:0033013) in L*er*, *tps1-11*, and *tps-12*. Seedlings were grown as described in the methods section. In brief, RNA was isolated from 14-day-old seedlings that were exposed to 37 °C for one hour or maintained at 22 °C as control ($N$ = 3). Library preparation and 100 bp single-end sequencing were performed by the Core Unit Systemmedizin at the University of Wuerzburg using the Illumina TruSeq RNA Library Prep Kit and an Illumina NextSeq2000 sequencer, respectively. Reads were aligned to the reference genome (TAIR10 release) using Bowtie2 and normalised to the total read number to obtain transcripts per million (TPM) values. Differentially expressed genes were detected using an ANOVA framework and an FDR-corrected $p$-value cut-off of 0.05 and filtered for genes with a $|\log_2(\text{fold change})| > 1$. GO term enrichment analysis was done using the PANTHER online tool and enriched GO terms were visualised using the ggplot2 package in *R* according to Bonnot *et al.*, (2019). DE: differentially expressed.
