## Supplementary material for "Natural variation of warm temperature-induced raffinose accumulation identifies *TREHALOSE-6-PHOSPHATE SYNTHASE 1* as a modulator of thermotolerance": Table S1

**Supplemental Table S1.** Multiple reaction monitoring (MRM) transitions and response factors used for the quantification of soluble carbohydrates.

***Analyte	MRM transition (m/z)	Response factor***

Fructose^a^	179 > 89	0.297

Glucose^a^	179 > 89	0.338

Inositol^a^	179 > 87	0.493

Galactinol^b^	341 > 179	1.157

Sucrose^b^	341 > 179	0.940

Raffinose^b^	503 > 179	0.458

Stachyose^b^	665 > 383	1.044

d_2_-Glucose (IS)	191 > 91	--

d_2_-Trehalose (IS)	343 > 180	--

^a^ These analytes were quantified relative to d2-glucose. ^b^ These analytes were quantified relative to d2-trehalose. IS: internal standard
