## Supplementary material for "Natural variation of warm temperature-induced raffinose accumulation identifies *TREHALOSE-6-PHOSPHATE SYNTHASE 1* as a modulator of thermotolerance": Table S2

**Supplemental Table S2.** Expression data of the genes within ± 5 kb of the single-nucleotide polymorphisms that were statistically significantly associated with the variation of warm temperature-induced raffinose accumulation at 32 °C.

| ***Locus identifier*** | ***Name*** | ***Transcripts per million^a^*** | | ***Log2FC*** | ***p-value^b^*** |
| --- | --- | --- | --- | --- | --- |
|  |  | ***22 °C*** | ***37 °C*** |  |  |
| **AT1G78540** | ***SHB*** | **2.9 (0.5)** | **1.9 (0.2)** | **-0.59** | **0.031** |
| **AT1G78550** | **na** | **5.1 (0.2)** | **3.4 (0.8)** | **-0.60** | **0.020** |
| **AT1G78560** | ***BASS1*** | **23.2 (0.9)** | **14.7 (1.4)** | **-0.65** | **0.001** |
| **AT1G78570** | ***RHM1*** | **121.0 (15.8)** | **37.0 (3.5)** | **-1.71** | **0.001** |
| **AT1G78580** | ***TPS1*** | **18.7 (2.4)** | **7.3 (1.1)** | **-1.35** | **0.002** |
| **AT1G78700** | ***BEH4*** | **22.3 (1.2)** | **8.2 (1.1)** | **-1.44** | **0.000** |
| AT1G78710 | na | 0.1 (0.1) | 0.0 (0.0) | na |  |
| AT1G78720 | na | 0.0 (0.0) | 0.0 (0.1) | na |  |
| **AT1G78730** | **na** | **2.4 (0.3)** | **1.4 (0.3)** | **-0.83** | **0.010** |
| AT1G78740 | na | 0.0 (0.0) | 0.0 (0.0) | na |  |
| **AT4G16990** | ***RLM3*** | **51.8 (7.0)** | **19.8 (1.9)** | **-1.39** | **0.002** |
| **AT4G17000** | **na** | **3.1 (0.6)** | **0.7 (0.1)** | **-2.20** | **0.002** |
| **AT5G18320** | ***PUB46*** | **0.2 (0.1)** | **0.3 (0.1)** | **0.19** | **0.678** |
| AT5G18330 | *PUB47* | 0.0 (0.0) | 0.1 (0.1) | na | 0.116 |
| **AT5G18340** | ***PUB48*** | **0.6 (0.4)** | **48.0 (6.9)** | **6.24** | **0.000** |
| AT5G18350 | na | 0.1 (0.1) | 0.0 (0.0) | na | 0.116 |
| **AT5G18360** | ***BAR1*** | **1.4 (0.2)** | **0.3 (0.1)** | **-2.19** | **0.001** |
| **AT5G18370** | ***DSC2*** | **1.7 (0.3)** | **0.4 (0.1)** | **-2.06** | **0.002** |

Genes that were considered candidate genes are set in bold. ^a^ Expression data was derived from an unpublished in-house data set. In brief, Columbia-0 (Col-0) seedlings were grown on ½-strength MS-Agar, pH 5.7, for twelve days under long-day conditions (16:8 h, light/dark, 22 °C) before they were subjected to a 37 °C heat treatment for one hour or maintained at 22 °C as control. Library preparation and sequencing were performed by BGI Genomics, Shenzhen, China. Data are given in transcripts per million and shown as means (SD). ^b^ Statistical significance was tested using a Student’s *t*-test.
