## Supplementary material for "Natural variation of warm temperature-induced raffinose accumulation identifies *TREHALOSE-6-PHOSPHATE SYNTHASE 1* as a modulator of thermotolerance": Table S3

**Supplemental Table S3.** Characterisation of the genetic backgrounds of *tps1* TILLING lines.

| ***Genotype*** | ***No. of SNPs vs. Col-0 reference*** | ***Relative composition*** | |  |
| --- | --- | --- | --- | --- |
|  |  | **L*er*** | **Col-0** | **other** |
| L*er* | 104 787 | 100% | -- | -- |
| *tps1-11* | 52 762 | 46.0% | 49.7% | 4.3% |
|  | 48 152 L*er*-type allele |  |  |  |
|  | 4 610 other accession |  |  |  |
| *tps1-12* | 67 987 | 58.0% | 37.8% | 4.2% |
|  | 63 411 L*er*-type allele |  |  |  |
|  | 4 576 other accession |  |  |  |

Transcriptome-wide detection of single-nucleotide polymorphisms (SNPs) in Landsberg *erecta* (L*er*), *tps1-11* and *tps1-12 versus* the Columbia-0 (Col-0) reference genome (TAIR10) using 100 bp single-end RNA-Seq data. Library preparation and sequencing were performed by the Core Unit Systemmedizin at the University of Wuerzburg using the Illumina TruSeq RNA Library Prep Kit and an Illumina NextSeq2000 sequencer, respectively. Note that detection of alleles other than L*er* or Col-0 in the TILLING lines might also be due to erroneous SNP calling or mistakes in the annotated reference genome.
